# Reduction of Glut1 in retinal neurons but not the RPE alleviates polyol accumulation and normalizes early characteristics of diabetic retinopathy

**DOI:** 10.1101/2020.08.31.275230

**Authors:** Nicholas C. Holoman, Jacob J. Aiello, Timothy D. Trobenter, Matthew J. Tarchick, Michael R. Kozlowski, Emily R. Makowski, Darryl C. De Vivo, Charandeep Singh, Jonathan E. Sears, Ivy S. Samuels

## Abstract

Hyperglycemia is a key determinant for development of diabetic retinopathy (DR). Inadequate glycemic control exacerbates retinopathy, while normalization of glucose levels delays its progression. In hyperglycemia, hexokinase is saturated and excess glucose is metabolized to sorbitol by aldose reductase via the polyol pathway. Therapies to reduce retinal polyol accumulation for the prevention of DR have been elusive due to low sorbitol dehydrogenase levels in the retina and inadequate inhibition of aldose reductase. Using systemic and conditional genetic inactivation, we targeted the primary facilitative glucose transporter in the retina, Glut1, as a preventative therapeutic in diabetic male and female mice. Unlike wildtype diabetics, diabetic *Glut1*^*+/−*^ mice did not display elevated Glut1 levels in the retina. Furthermore, diabetic *Glut1*^*+/−*^ mice exhibited ameliorated ERG defects, inflammation and oxidative stress, which was correlated with a significant reduction in retinal sorbitol accumulation. RPE-specific reduction of Glut1 did not prevent an increase in retinal sorbitol content or early hallmarks of DR. However, like diabetic *Glut1*^*+/−*^ mice, reduction of Glut1 specifically in retinal neurons mitigated polyol accumulation and completely prevented retinal dysfunction and the elevation of markers for oxidative stress and inflammation associated with diabetes. These results suggest that modulation of retinal polyol accumulation via Glut1 in photoreceptors can circumvent the difficulties in regulating systemic glucose metabolism and be exploited to prevent DR.

**Significance:** Diabetic retinopathy (DR) affects one third of diabetic patients and is the primary cause of vision loss in adults aged 20-74. While anti-VEGF and photocoagulation treatments for the late-stage vision threatening complications can prevent vision loss, a significant proportion of patients do not respond to anti-VEGF therapies and mechanisms to stop progression of early-stage symptoms remain elusive. Glut1 is the primary facilitative glucose transporter for the retina. We determined that a moderate reduction in Glut1 levels, specifically in retinal neurons, but not the RPE, was sufficient to prevent retinal polyol accumulation and the earliest functional defects to be identified in the diabetic retina. Our study defines modulation of Glut1 in retinal neurons as a targetable molecule for prevention of DR.

## Introduction

Hyperglycemia is a primary risk factor for the development of diabetic retinopathy (DR) (Lee et al., 2015; Lima et al., 2016; Sabanayagam et al., 2016). Increased glucose metabolism and retinal polyol accumulation are key pathological features of DR (Gabbay, 1973; Asnaghi et al., 2003; Dagher et al., 2004) and directly contribute to DR via the generation of reactive oxygen species and advanced glycation end products, a reduction in pools of reduced glutathione, and increased retinal osmolarity (Lorenzi, 2007). However, successful therapies targeting glucose and polyol breakdown have been elusive. Inhibition of the two key metabolic enzymes in the polyol pathway, aldose reductase and sorbitol dehydrogenase, has not been possible due to an inability to find a balance between efficacy and tolerance with currently available therapeutics. As altered retinal function (Aung et al., 2013; Samuels et al., 2015), increased oxidative stress and inflammation (Du et al., 2003; Al-Kharashi, 2018) and neurodegeneration (van Dijk et al., 2012; Sohn et al., 2016), are each found at early time points of hyperglycemia (Robinson et al., 2012) and are refractive to reductions in retinal glucose and polyols (Obrosova et al., 2003; Sun et al., 2006), we sought to identify an alternative mechanism to inhibit polyol accumulation and prevent DR.

Glut1 (encoded by *Slc2a1*) is the primary facilitative glucose transporter for the retina/RPE (Rizzolo, 1997). It is localized to both the apical and basal membranes of the RPE, and throughout the retina, including rod and cone photoreceptors, retinal ganglion cells and Müller glia (Kumagai et al., 1994). Previous studies demonstrate that reduction of *Slc2a1* levels in the retina with siRNA (Lu et al., 2013; You et al., 2017) or pharmacological inhibition of Glut1 (You et al., 2018) decreased retinal pathophysiology in the streptozotocin (STZ) mouse model of diabetes. However, these studies did not identify the key molecular components involved in mediating this effect.

We previously correlated the time course and extent of ERG defects in STZ-induced diabetic mice with hyperglycemia (Samuels et al., 2015). Reduced ERG amplitudes and increased ERG latencies occur prior to structural changes to the retina and are predictive of microaneurysm development and DR severity (Ng et al., 2008; Ratra et al., 2020). Molecular targets that prevent ERG defects could be utilized for preventative or interventional therapies. Herein, we first investigated whether *Slc2a1* expression and/or Glut1 protein levels in the retina and RPE were significantly different at early DR stages that exhibit ERG defects. We report that DR is associated with elevated retinal Glut1 levels, without changes in expression of *Slc2a1* or other glucose transporters. We next used a genetic approach to reduce Glut1 and found that systemic Glut1 haploinsufficiency in *Glut1*^*+/−*^ mice protected against DR phenotypes including altered electroretinography, polyol accumulation and increased retinal oxidative stress and inflammation. The protection was retina-specific, as reduction of Glut1 in retinal neurons conferred a similar prevention of DR while reduction of Glut1 in the RPE did not. These data demonstrate that although the RPE serves to supply the retina with glucose for proper retinal homeostasis, manipulation of Glut1 levels in the retina, but not the RPE, is a valuable target for therapies to prevent and treat DR. Moreover, reduction of retinal sorbitol and prevention of DR can be achieved by modulation of Glut1 rather than manipulation of key enzymes in glucose metabolism.

## Materials and Methods

### Ethical Approval

Treatment of animals followed the ARVO Resolution on Treatment of Animals in Research, and all animal procedures were approved by the Institutional Animal Care and Use Committee of the Louis Stokes Cleveland VA Medical Center.

### Mice

*Glut1*^*+/−*^ mice were kindly provided by Darryl de Vivo (Columbia University), *VMD2*^*Cre/+*^ mice by Joshua Dunaief (University of Pennsylvania), *Crx*^*Cre/+*^ mice by Sujata Rao (Cleveland Clinic; currently available from RIKEN BRC #RBRC05426) and *Glut1*^*flox*^ mice by E. Dale Abel (University of Iowa; currently available from The Jackson Laboratory #031871). C57Bl/6J were purchased from The Jackson Laboratory (#000664). At 6-8 weeks of age, in both male and female mice, diabetes was induced by three sequential daily intraperitoneal injections of a freshly prepared solution of STZ in 0.1 M citrate buffer (pH 4.4) at 30 mg/kg body weight. In the STZ group, insulin (0–0.2 units of neutral protamine Hagedorn (NPH) Humulin N, Eli Lilly and Co., Indianapolis, IN) was given by intraperitoneal injection every other day, as needed post hyperglycemia, to prevent ketosis without preventing hyperglycemia and glucosuria. CNTL mice received citrate buffer only and did not receive insulin.

### Experimental Design and Statistical Analysis

For all analyses, data were compiled as mean ±SEM or SD as indicated in figure legends, and statistics were performed on GraphPad Prism 6 using non-repeated measures, one-way or two-way ANOVA with Tukey post-hoc analysis (GraphPad, Inc., La Jolla, CA, USA). Statistical significance was determined by achieving a p value for both the ANOVA and multiple comparisons test below 0.05. At least three animals per condition per time point were used for all experiments. Full details for each experiment including group numbers, statistical tests and test values are included in Table 1.

**Table 1:**
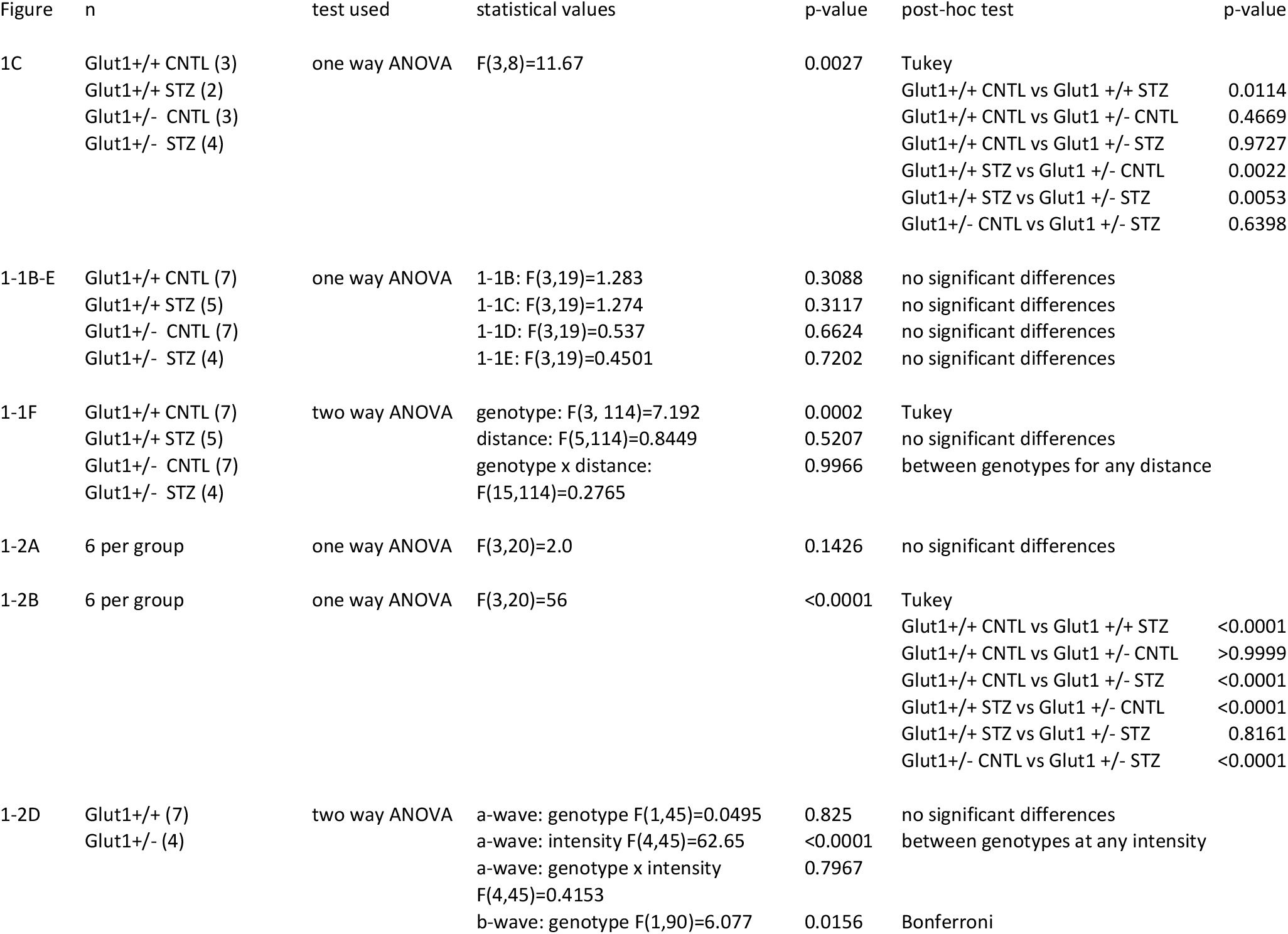
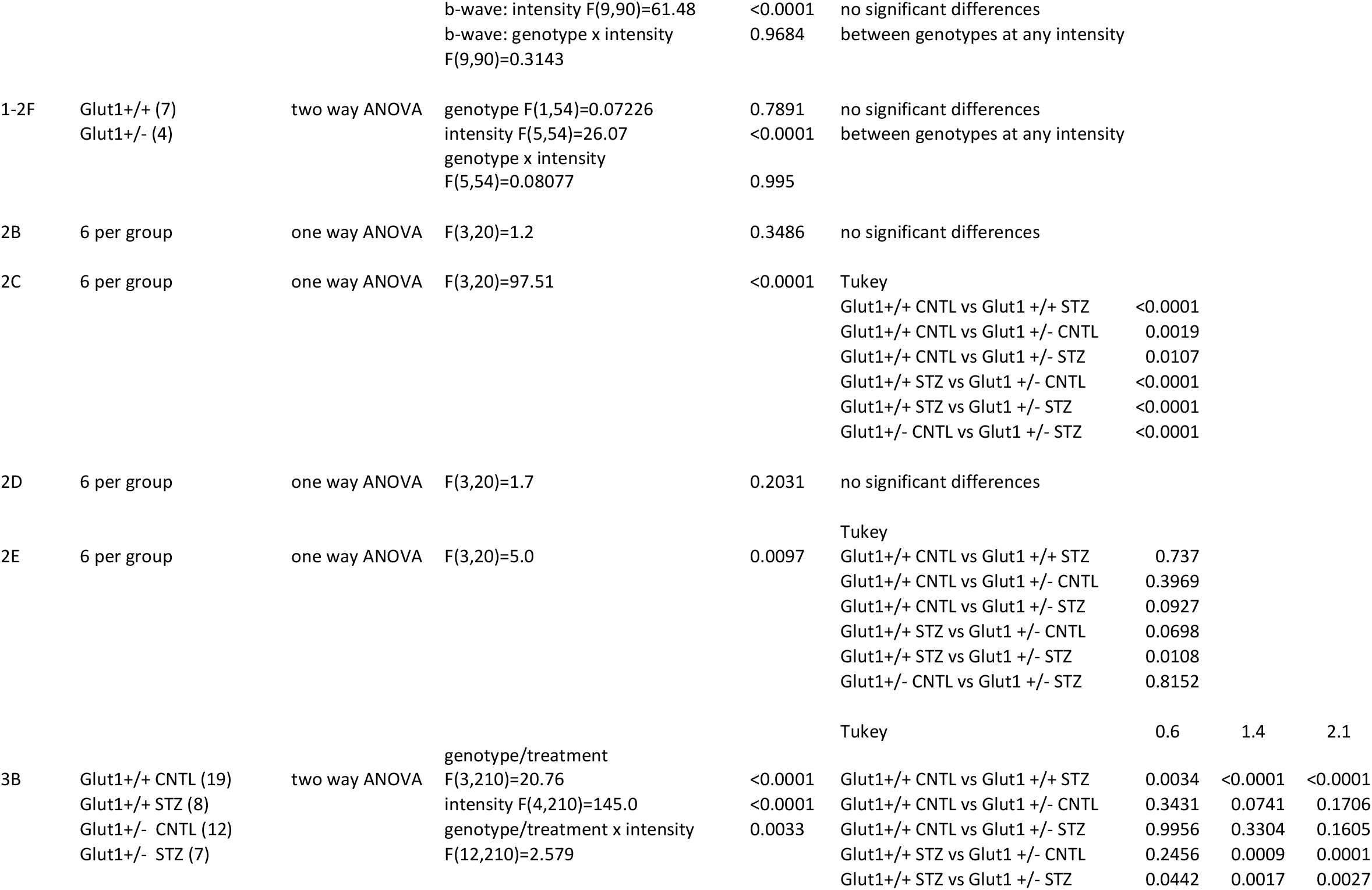
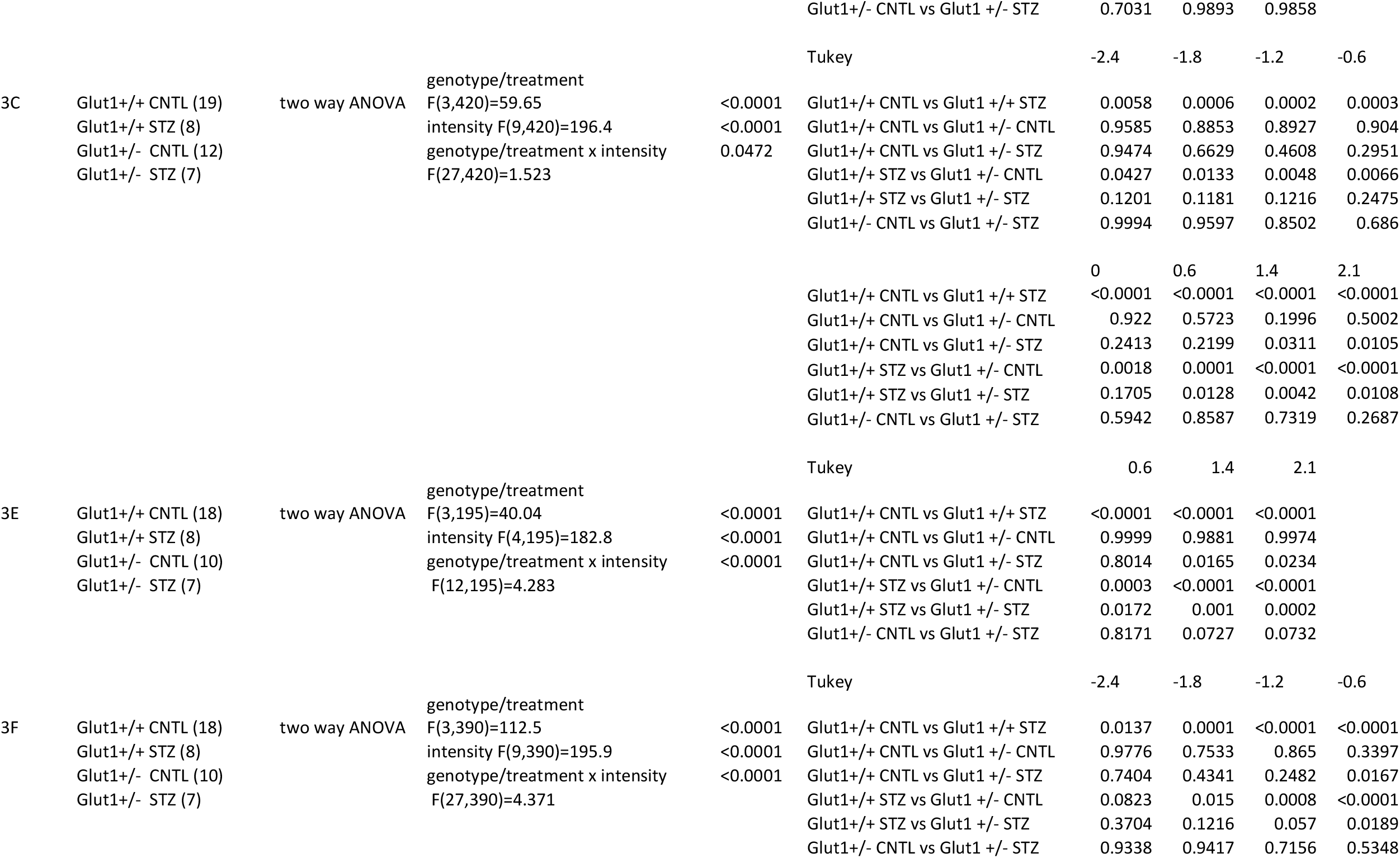
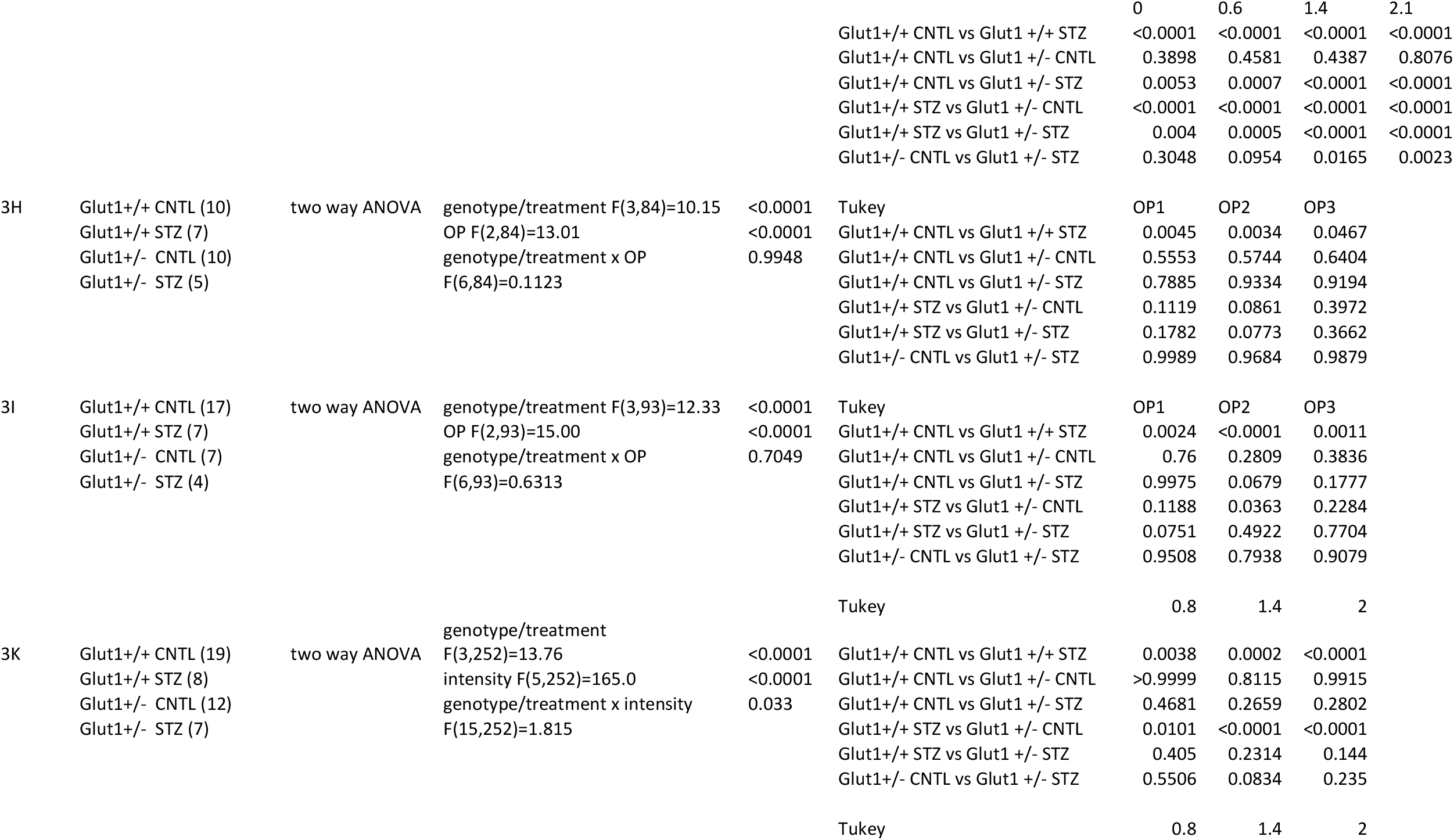
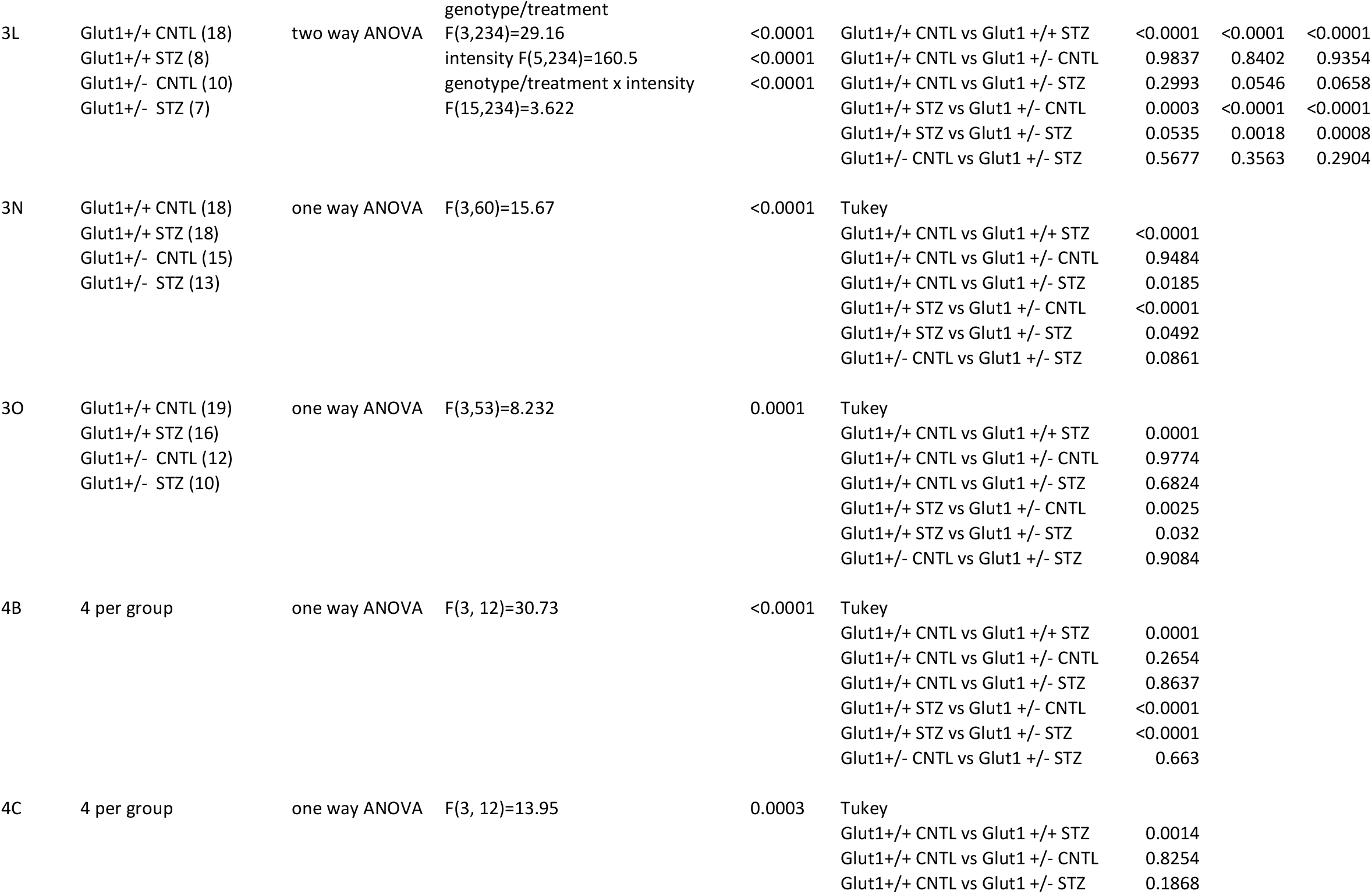
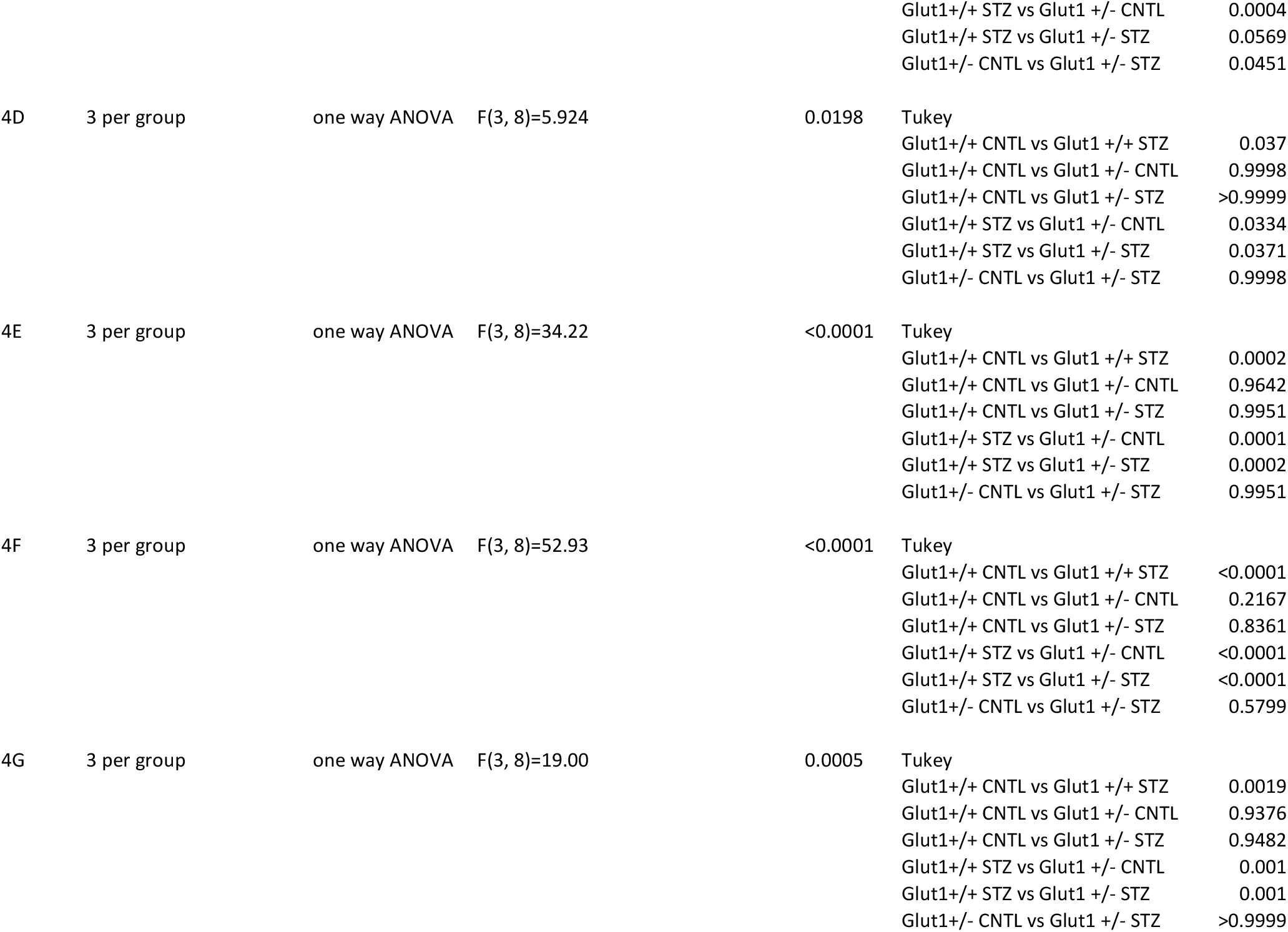
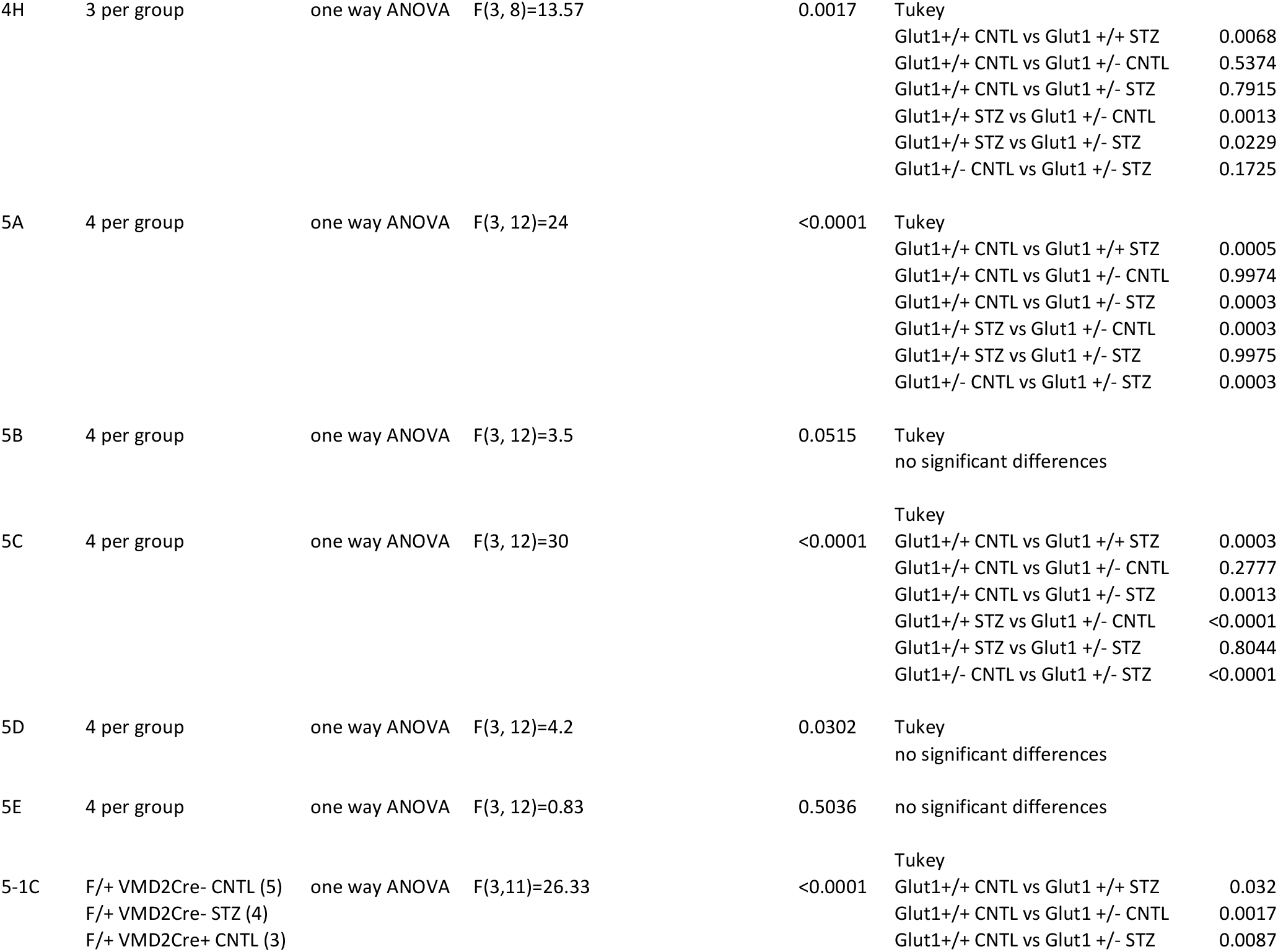
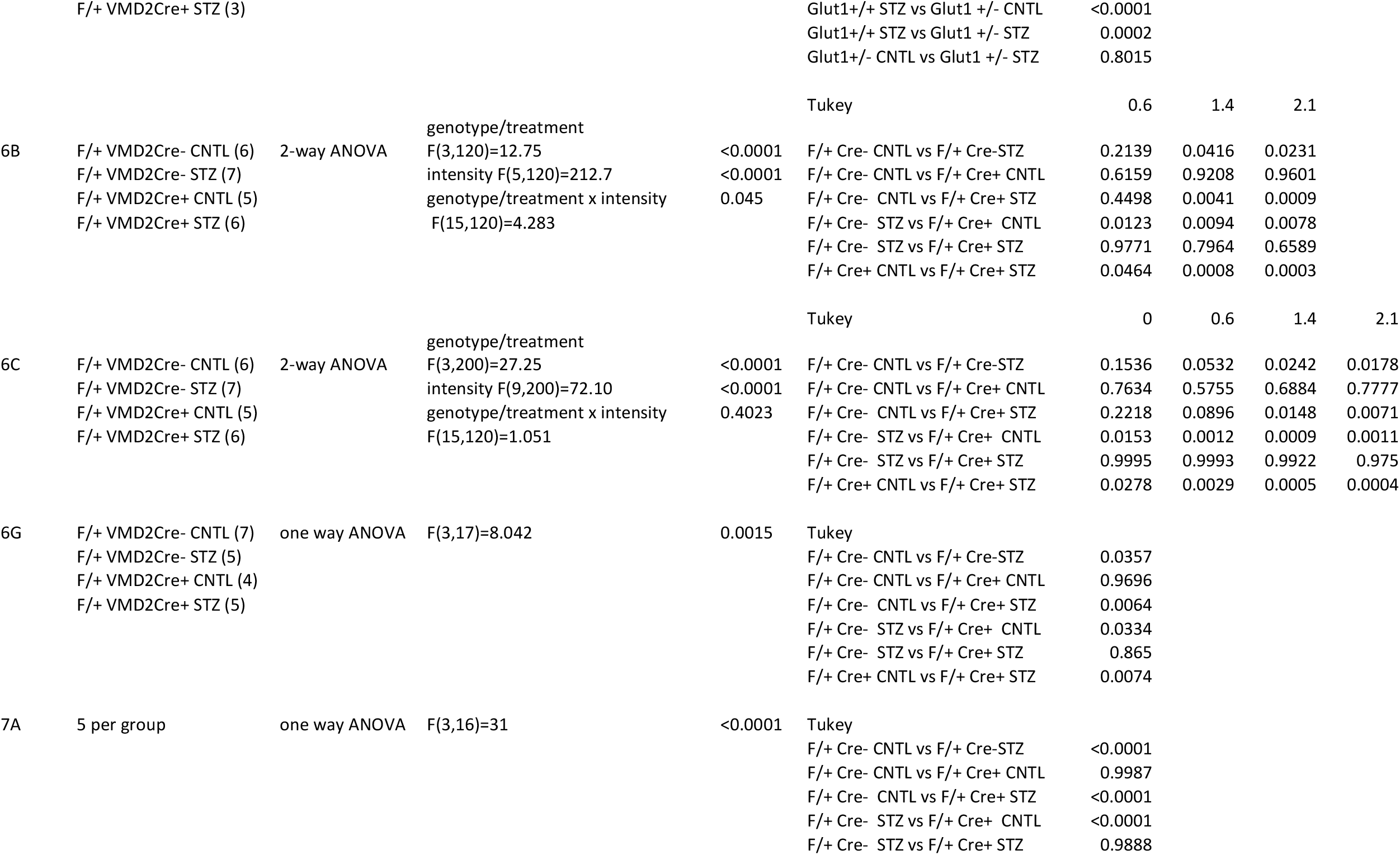
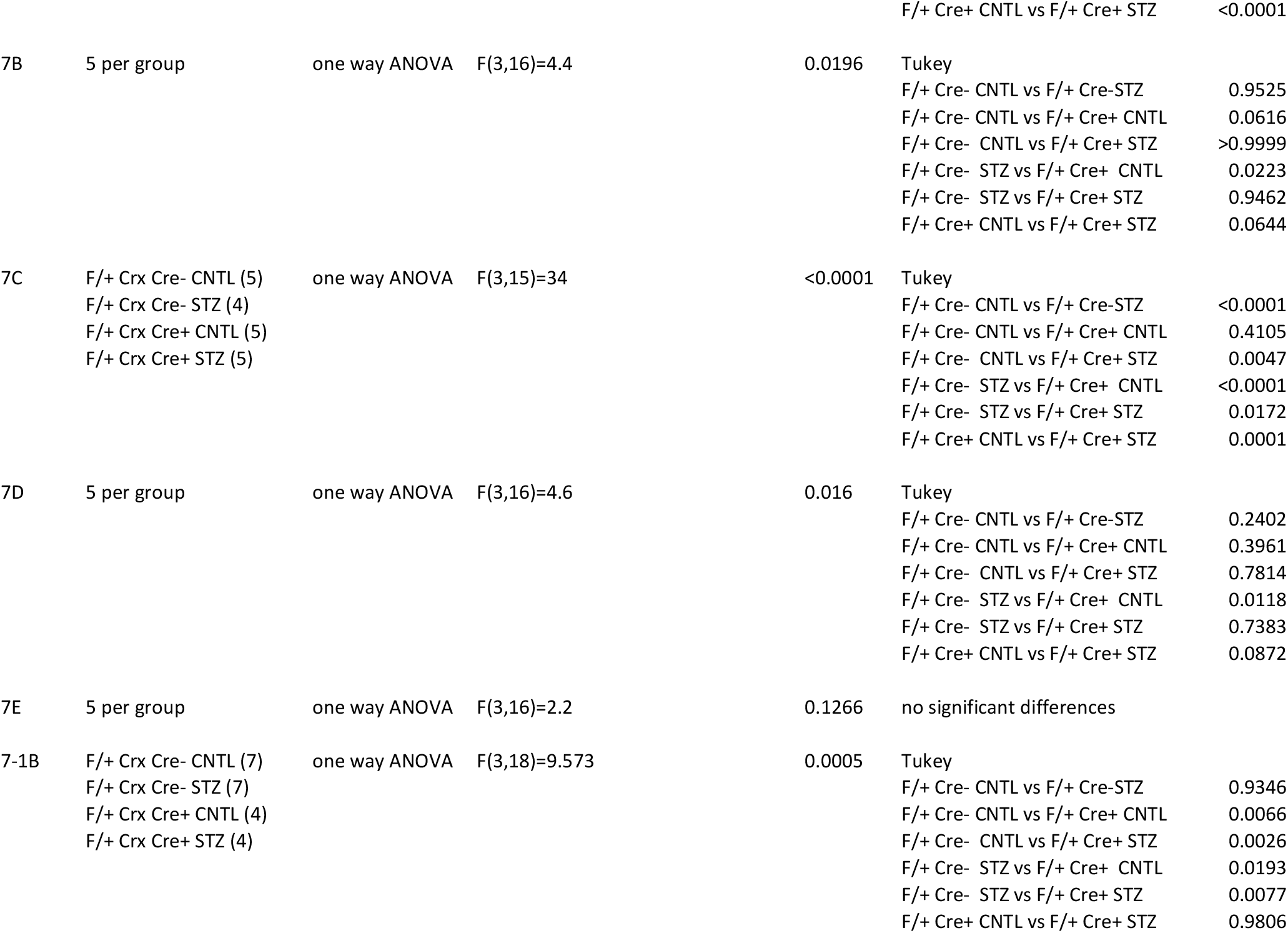
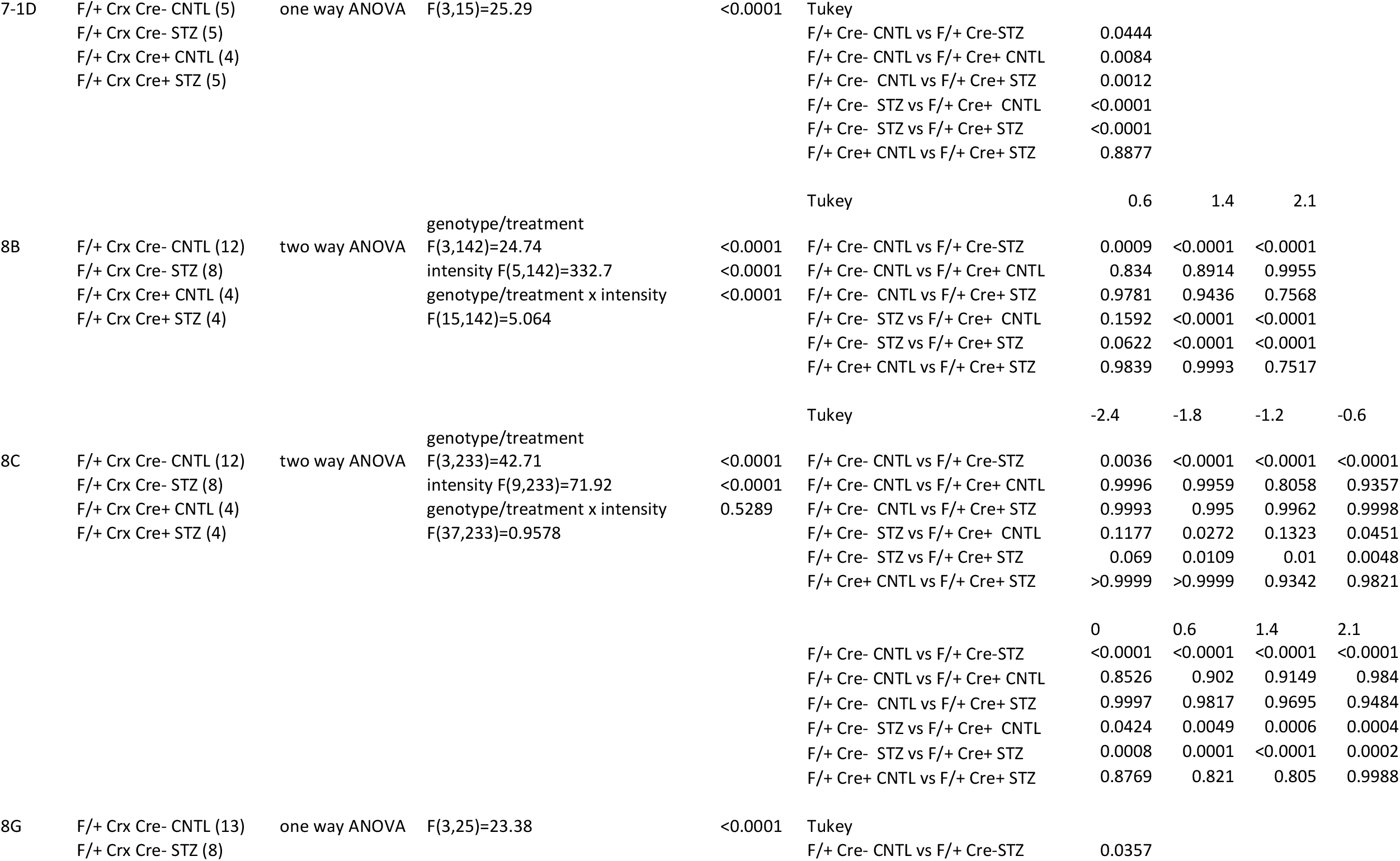
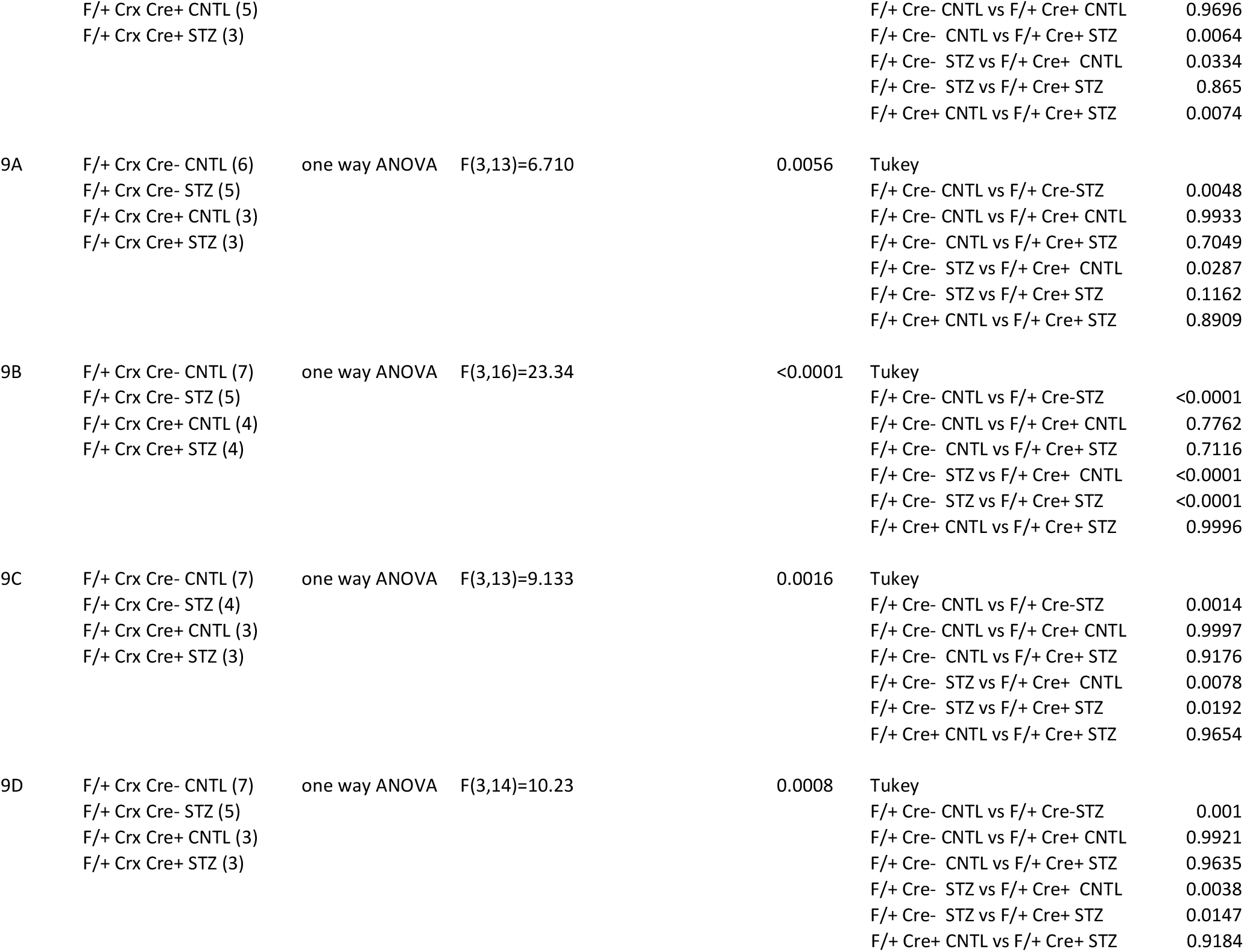
Experimental Design and Statistics

### Genotyping

The *Slc2a1 a*llele was identified by genotyping with the following primers:

SF3 5’-CCA TAA AGT CAG AAA TGG AGG GAG GTG GTG GT-3’

E1R 5’-GCG AGA CGG AGA ACG GAC GCG CTG TAA CTA-3’

NR 5’-CTA CCG GTG GAT GTG GAA TGT GTG CGA GGC-3’

The floxed *Slc2a1* allele was identified by genotyping with the following primers:

FRT-F 5’-CTC CAT TCT CCA AAC TAG GAA C-3’

FRT-R2 5’-GAA GGC ACA TAT GAA ACA ATG-3’

2.85F 5’-CTG TGA GTT CCT GAG ACC CTG-3’

2.9R 5’-CCC AGG CAA GGA AGT AGT TC-3’

The presence of Cre recombinase was identified by genotyping with the following primers:

CreF 5’-TGC CAC GAC CAA GTG ACA GCA ATG-3’

CreR 5’-ACC AGA GAC GGA AAT CCA TCG CTC-3’

### Electroretinography

After overnight dark adaptation, mice were anesthetized with 65mg/kg sodium pentobarbitol. Eye drops were used to anesthetize the cornea (1% proparacaine HCl) and to dilate the pupil (2.5% phenylephrine HCl, 1% tropicamide, and 1% cyclopentolate HCl). Mice were placed on a temperature-regulated heating pad throughout the recording session which was performed as previously described (Samuels et al., 2015). Amplitude of the a-wave was measured at 8.3 ms following the stimulus. Amplitude of the b-wave was calculated by summing the amplitude of the a-wave at 8.3ms with the peak of the waveform after the oscillatory potentials (≥40ms). Light-adapted response amplitudes were calculated by summing the peak of the waveform with the amplitude at 8.3ms. OP amplitude was determined by measuring the change in amplitude from the preceding trough to the peak of each potential. Amplitude of the c-wave was determined by subtracting the average baseline amplitude from the maximal response following the b-wave.

### Histology and Light Microscopy

Enucleated eyes were fixed in 0.1 M sodium cacodylate buffer (pH 7.4) containing 2% formaldehyde and 2.5% glutaraldehyde. The tissues were then osmicated, dehydrated though a graded ethanol series, plasticized in acetonitrile, and embedded in epoxy resin (Embed-812/DER73 Epon kit; Electron Microscope Services, Hatfield, PA, USA). Semi-thin sections (0.8 µm) were cut along the horizontal meridian through the optic nerve and stained with 1% toluidine blue O for evaluation. Photomicrographs were taken of sections traversing the optic nerve. The distance from the outer limiting membrane (OLM) to the inner limiting membrane (ILM) was measured in three sections per animal and averaged for a minimum of three animals per group using ImageJ software. Additional photomicrographs were taken 250 µm from the optic nerve, and the length of the outer segment (OS), inner segment (IS), and outer nuclear layer (ONL) were measured in three equidistant areas per section per animal. Thickness of RPE was also measured at 3100x magnification from three sections of each mouse.

### Immunohistochemistry

After mice were euthanized and enucleated, eyes were fixed in 0.1 M sodium phosphate buffer (pH 7.4) containing 4% paraformaldehyde. After removal of the cornea and lens, the posterior pole was immersed through a graded series of sucrose solutions as follows: 10% for 1 h, 20% for 1 h, and 30% overnight. Eyes were embedded in OCT freezing medium, flash frozen on powderized dry ice, and immediately transferred to −80°C. Tissue was sectioned at 10 µm thickness at −30°C, mounted on superfrost slides, and stored at −80°C until processed. Sections were blocked in 0.1% Triton X-100, 1% bovine serum albumin, and 5% normal goat serum in phosphate-buffered saline (PBS) for 1 h at room temperature (RT) and then washed three times with PBS for 5 min each time. The sections were incubated overnight at 4°C with the primary antibody. Sections were rinsed with PBS three times for 10 min each time and incubated with secondary antibody (Alexa 488 or Alexa 594, 1:500; Molecular Probes) for 1 h at RT. After rinsing sections three times for 10 min each time with PBS, sections were mounted with DAPI (1:10,000 in 50% glycerol:PBS). Primary antibodies used were rabbit anti-Glut1 (Millipore #07-1401, 1:500), mouse anti-Glut1 (Abcam #ab40084, 1:100), rabbit anti-recoverin (Millipore #ab5585, 1:1000). Imaging was performed using a Leica laser scanning confocal microscope (TCSSP2, Leica Microsystems).

### Western blotting

Retinas were lysed on ice for 10 min in lysis buffer (20mM HEPES, 150mM NaCl, 1.5mM NgCl2, 2mM EGTA with 0.5% TritonX-100) containing protease inhibitors (Roche #5892970001) and phosphatase inhibitors (10mM NaF, 1mM PMSF, 1mM Na_3_(VO_3_)_4_, 12.5mM β-glycerophosphate and 2mM DTT) followed by sonication. Protein concentration was determined by 660nm BCA Assay (ThermoScientific #22660) and equivalent amounts of reduced protein (6x Laemelli sample buffer with 5% betamercaptoethanol) were separated by SDS-PAGE on 4-15% acrylamide gels. Proteins were transferred to PVDF membranes which were blocked with Intercept blocking buffer and imaged using IRDye 800CW goat anti-rabbit and 680 goat anti-mouse secondary antibodies (Li-Cor Biosciences, 1:10,000). Primary antibodies used included: Rb anti-Glut1 (Millipore #07-1401, 1:2000) and Mo anti-actin (Cell signaling #3700S, 1:1000). PVDF membranes were scanned with an Odyssey infrared scanner and densitometry was performed using LiCor Image Studio Software.

### Oxidative stress

Reactive oxygen species was measured in retinal cryosections by dihydroethidium (DHE) staining. Eyes were dissected on ice cold PBS and frozen in OCT embedding buffer within 15 minutes of enucleation. Fresh frozen retinal cryosections spanning the optic nerve were incubated with DHE (ThermoFisher, 1:5000) for 20 minutes followed by counterstaining with DAPI (1:10,000). Quantification of reactive oxygen species was calculated using NIH ImageJ software.

### Quantitative PCR

Gene expression of glucose transporters, inflammatory cytokines and oxidative stress molecules was measured by quantitative PCR on dissected retinal tissue. RNA was extracted using the RNAeasy Mini Kit (Quiagen #74104) and RT-PCR was performed using the Verso cDNA synthesis kit (ThermoScientific #AB1453A). Radiant Green 2X qPCR Lo-ROX enzyme was used for all qPCR. Relative fold changes in gene expression were determined using the comparative Ct method (2ΔΔCt method). Actin or 18S was used as the reference gene. HIF-1α (#QT01039542) and VEGF (#QT00160769) were analyzed using primers from Qiagen. Primers for all other genes investigated are listed in the table below:

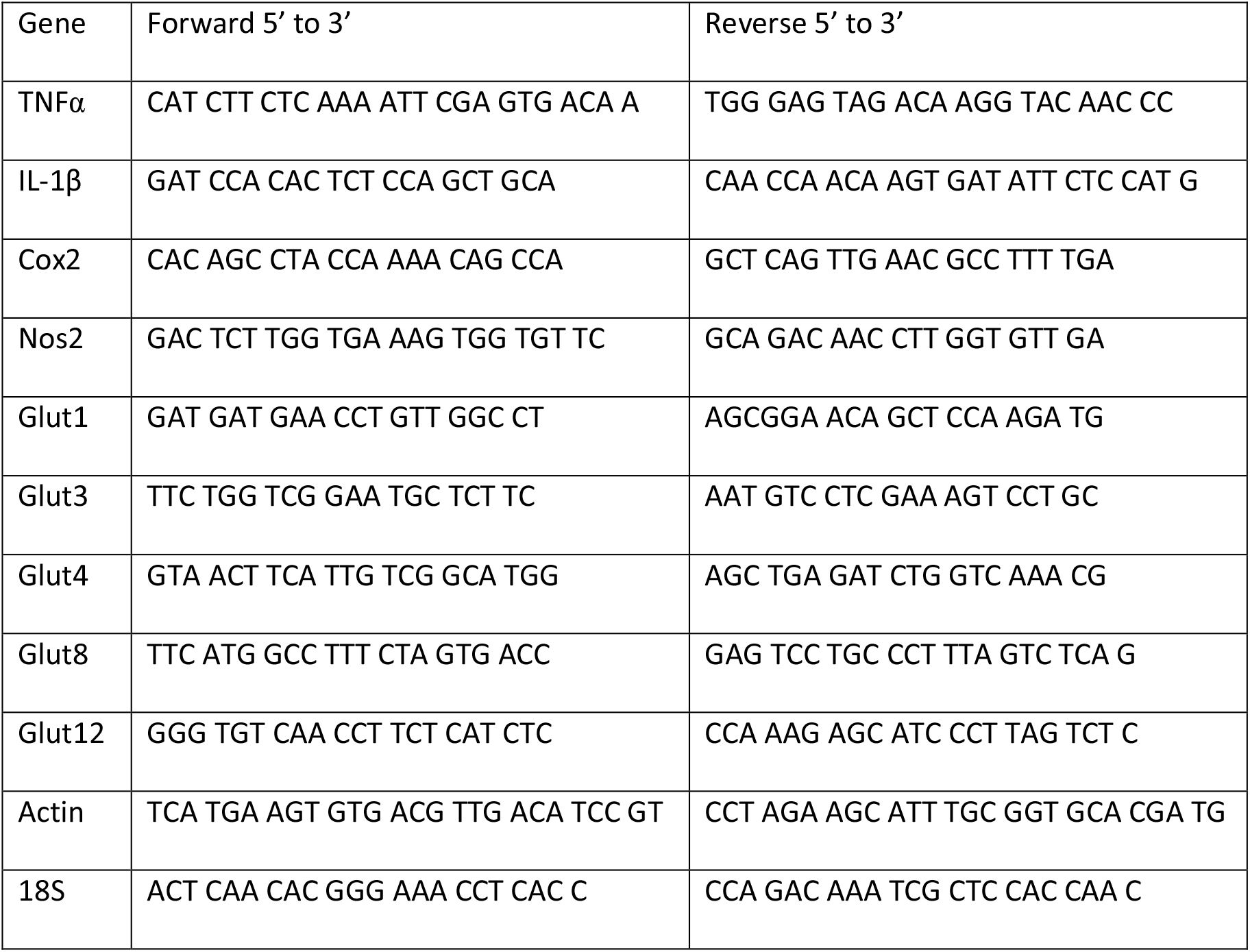

### Gas Chromatograpy-Mass Spectrometry

As previously described (Singh et al., 2020), for each mouse, both retinas were dissected on ice cold HBSS and flash frozen. To each tube, 500 µl of −20°C 80% methanol was added with 20 µl of 0.05 mg/ml of [^13^C_5_] ribitol as internal standard. Metabolites were extracted by sonication. Samples were then centrifuged at 15000 x g for 5 min at 4°C. After centrifugation, 300 µl of supernatant was transferred to fresh tube. Samples were dried overnight in a −4°C vacuum evaporator. Dried samples were first derivatized by adding 25 µl of 40 mg/ml methoxyamine in pyridine and then incubating on a thermomixer at 45°C for 30 min with 1000 rpm shaking speed. Samples were then additionally derivatized by adding 75ul of MSTFA + 1% TMCS and incubating on thermomixer as in the first step. One microliter of each sample was injected into the 7890B GC connected to 5977 MSD Agilent GCMS system. Injections were made in splitless or split 15 mode. GC column used was DB-5 ms 30 m × 0.25 mm × 0.25 µm with DuraGuard 10 m. Front inlet was set at 250°C with septum purge flow of 3 ml/min of helium. Samples were analyzed in a constant flow mode with helium set to 1.1 ml/min. GC method was 1 min at 60°C, followed by 10 °C/min increments until 325 °C and finally held at 325 °C for 10 min. Metabolites were measured in full scan mode using electron ionization with a scan window from 50 to 800 m/z. Solvent delay of 6.6 minutes was applied.

## Results

### Hyperglycemia-induced elevations in retinal Glut1 are not found in diabetic *Glut1*^*+/−*^ mice

It is widely accepted that functional defects in the light evoked responses of the retina occur in rodent models of diabetes (Aung et al., 2013; Samuels et al., 2015) and in diabetic patients (Greenstein et al., 1993; Tyrberg et al., 2011; Bearse and Ozawa, 2014; Ratra et al., 2020), both of which also demonstrate increased polyol accumulation (Gabbay, 1973; Dagher et al., 2004; Lorenzi, 2007). In the STZ model of diabetes, reductions in ERG amplitudes were correlated with hyperglycemia, at both 2 weeks and 4 weeks of diabetes (Samuels et al., 2015). Due to the association between onset of ERG defects and hyperglycemia, we first investigated whether Glut1 levels differed between diabetic and non-diabetic mice at these early time points. Confocal imaging demonstrated that in STZ-induced diabetic mice, Glut1 levels were increased throughout the retina, and notably in the inner segments and outer nuclear layer in comparison with the non-diabetic controls (Fig 1A). Based on the premise that acute reduction of Glut1 in the retina (*Slc2a1*) via siRNA injection or pharmaceutical inhibition (forskolin) reduced hallmarks of DR (Lu et al., 2013; You et al., 2017; You et al., 2018), we hypothesized that early characteristics of DR would be ameliorated in diabetic *Glut1*^*+/−*^ mice which exhibit a genetic, systemic 50% reduction in Glut1 by virtue of expressing only one *Slc2a1* allele. Non-diabetic *Glut1*^*+/+*^ and *Glut1*^*+/−*^ mice display indistinguishable morphology (Fig 1-1), similar body weights (Fig 1-2A) and identical light-evoked responses of the retina (electroretinography; Fig 1-2C-F). *Glut1*^*+/+*^ and *Glut1*^*+/−*^ mice also exhibit comparable STZ-induced increases in blood glucose levels (Fig 1-2B). Confocal microscopy of retinal cryosections from *Glut1*^*+/+*^ and *Glut1*^*+/−*^ mice stained with anti-Glut1 (red) and anti-Recoverin (green, photoreceptor marker) antibodies confirmed that 4wk diabetic (STZ) *Glut1*^*+/+*^ mice exhibited elevated Glut1 in the retina and RPE in comparison with non-diabetic (CNTL) *Glut1*^*+/+*^ mice (Fig 1B). As expected, CNTL *Glut1*^*+/−*^ retinas displayed significantly lower Glut1 levels in the retina and RPE than the *Glut1*^*+/+*^ mice. Additionally, while Glut1 was upregulated in *Glut1*^*+/+*^ STZ mice, a similar magnitude increase was not observed in *Glut1*^*+/−*^ STZ retinas. Quantification of retinal Glut1 levels by western blot analysis confirmed a 2.15-fold increase of retinal Glut1 in wildtype diabetics (Fig 1C-D). However, no significant difference was found in retinal Glut1 levels between diabetic and non-diabetic *Glut1*^*+/−*^ mice, and diabetic *Glut1*^*+/−*^ mice had 0.4-fold lower Glut1 levels compared to diabetic *Glut1*^*+/+*^ mice (Fig 1D; F(3,8)=11.67, p=0.0027).

**Figure 1:**
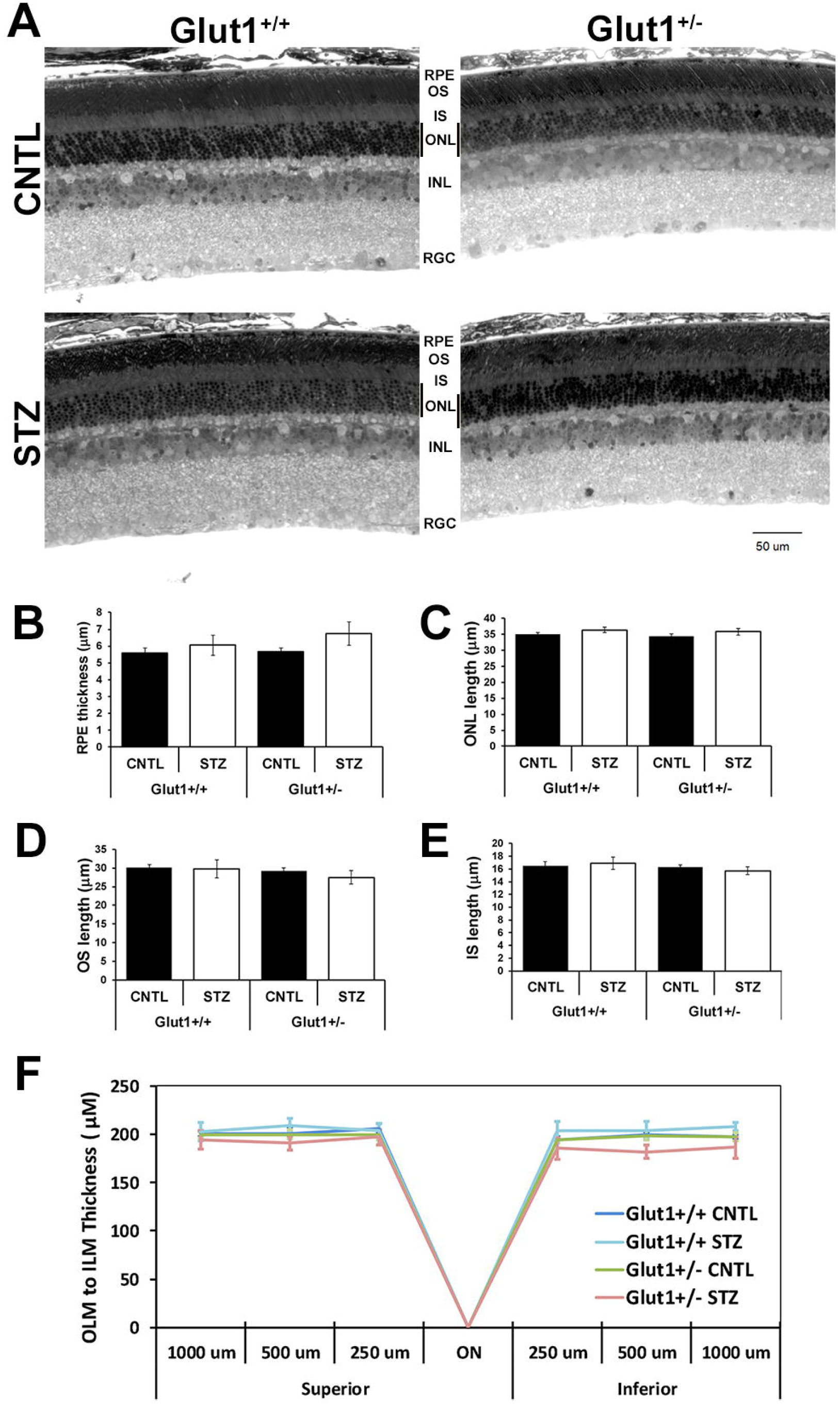
Retinal Glut1 expression and protein levels are not elevated in diabetic *Glut1*^*+/−*^ mice. A. Glut1 immunoreactivity (red) in confocal images taken from wildtype control and STZ-injected mice following 4 weeks of diabetes. Scale bar= 50µm. RPE, retinal pigmented epithelium; OS, outer segments; IS, inner segments; ONL, outer nuclear layer; OPL, outer plexiform layer; INL, inner nuclear layer; IPL, inner plexiform layer; RGC, retinal ganglion cell layer. B. Confocal images of Glut1 and recoverin immunoreactivity in cryosections following 4 weeks of diabetes. Scale bar=50µm. C. Protein levels of Glut1 from dissected retinas following 4 weeks of diabetes. Retinas were dissected and total Glut1 levels were normalized to β-actin for quantitative analysis. The graph depicts mean ± SEM. n≥3 in each group. *p≤0.05; **p≤0.001; ***p≤0.0001. Extended Figure 1-1 depicts normal retinal morphology in *Glut1*^*+/−*^ mice and Figure 1-2 illustrates their normal electroretinography and responses to diabetes. Table 1-1 presents real-time PCR of glucose transporter expression in the retina of each cohort of mice.

Analysis of mRNA expression of Glut1 in the retina of each cohort of mice verified a reduction in *Slc2a1* expression in *Glut1*^*+/−*^ mice as compared to *Glut1*^*+/+*^ littermates (Table 1-1; F(3,10)=8.898, p=0.0035). No significant differences in *Slc2a1* were found between control and diabetic *Glut1*^*+/−*^ mice. And as previously reported (Fernandes et al., 2004), no significant change in *Slc2a1* mRNA expression was found as a result of diabetes (Table 1-1, *Glut1*^*+/+*^ CNTL vs STZ and *Glut1*^*+/−*^ CNTL vs STZ). Analysis of other retinal glucose transporters demonstrated that there was also no significant change in expression of Glut3/*Slc2a3*, Glut4/*Slc2a4*, Glut8/*Slc2a8* or Glut12/*Slc2a12* in the retina of each cohort of mice after 4 weeks of diabetes (Table 1-1). These data illustrate that diabetes does not affect mRNA expression of glucose transporters in the retina, and that reduction of *Slc2a1* levels was not compensated for by an increase in expression of other glucose transporters.

### Reduction of Glut1 normalizes polyol accumulation, retinal dysfunction and increased inflammation/oxidative stress

Since systemic reduction of Glut1 normalized retinal Glut1 levels in diabetic mice to non-diabetic levels, we sought to determine if glucose transport and metabolism in the retina was modulated. Concentration of retinal glucose and glucose metabolites were measured by GC/MS in mice fasted for ≥7 hours. Although overt systemic hyperglycemia was observed by analysis of blood glucose levels (Fig 1-2A; F(3,20)=56, p<0.0001), no difference in retinal glucose levels between genotypes or treatment was identified (Fig 2A; F(3,20)=1.2, p=0.3486). A significant increase in retinal sorbitol was identified in diabetic *Glut1*^*+/+*^mice, which was significantly mitigated in diabetic *Glut1*^*+/−*^ mice (Fig 2B; F(3,20)=98, p<0.0001). Sorbitol and mannitol are isomers and have very similar EI mass spectra. Using our GCMS method, we were able to discern the identity of the polyol peaks by chromatographic separation (Fig 2-1) to verify the increase in sorbitol. Interestingly, while there was no change in retinal fructose levels (Fig 2C; F(3,20)=1.7, p=0.2031), the lactate:pyruvate ratio was slightly higher in the wildtype diabetic mice compared to *Glut1*^*+/−*^ diabetics, but not any other group (Fig 2D; F(3,20)=5.0,p=0.0097). These results demonstrate that the polyol branch of glucose metabolism had normal metabolite levels in *Glut1*^*+/−*^ retinas and that reduction of Glut1 prevented retinal polyol accumulation.

**Figure 2:**
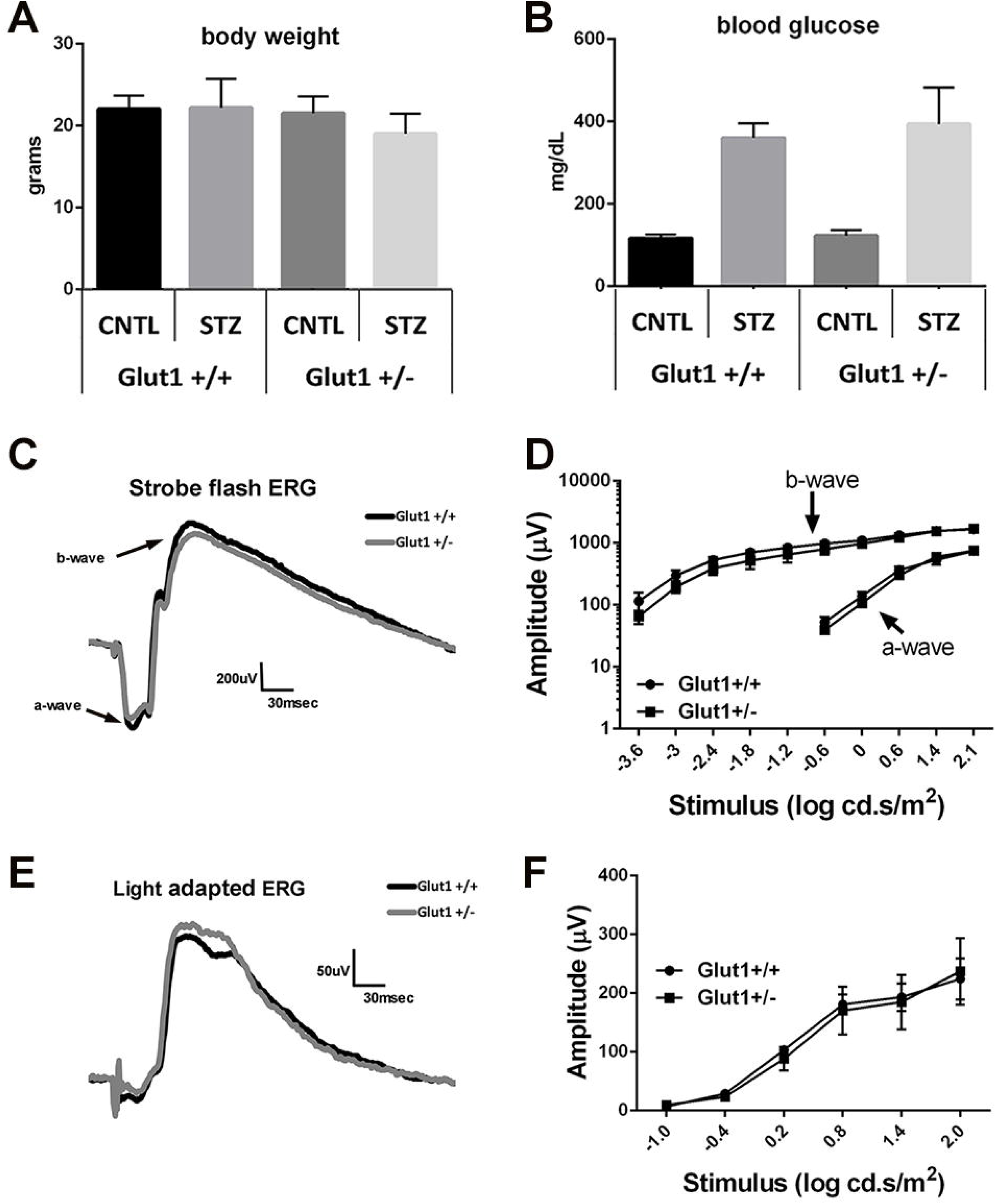
Systemic reduction of Glut1 in diabetic mice reduces retinal polyol accumulation. A. Glucose is metabolized to sorbitol by aldose reductase (AR) which is abundantly expressed in the retina. Sorbitol catabolism to fructose occurs via sorbitol dehydrogenase (Sord), which is present in extremely low levels in the retina. B-D. GC/MS was utilized to perform metabolomics on retinas from fasted mice at 4 weeks of diabetes. Relative quantities of glucose (B), sorbitol (C) and fructose (D) were normalized to ^13^C_5_-ribitol for comparison between genotypes. E. The lactate:pyruvate ratio was calculated as a surrogate for measurement of cytosolic NADH/NAD^+^. Graphs represent mean ± SD. n=6 for each group. *p≤0.05; **p≤0.001; ***p≤0.0001. Figure 2-1 shows the extracted ion chromatogram m/z 319 of mannitol and sorbitol authentic standards, demonstrating the baseline separation of these compounds on the GC column.

To determine if the normalization of retinal sorbitol content correlated with physiology, ERGs were measured after 2 and 4 weeks of diabetes. ERG defects are frequently observed prior to cell loss or the development of structural changes in the retina. Therefore, we chose to perform the ERG at both 2 and 4 weeks to identify the earliest time point of functional alterations. *Glut1*^*+/−*^ mice displayed normal ERG waveforms at baseline (6-8 weeks of age, before STZ injections; Figure 1-1C-E) and no differences were observed between non-diabetic *Glut1*^*+/+*^ and *Glut1*^*+/−*^ mice at 2 or 4 weeks post saline injections (Fig 3). Vision is also clinically spared in patients with Glut1 deficiency syndrome. In line with our previous findings (Samuels et al., 2015), significant reductions in both a- and b-wave amplitudes were present across all stimulus intensities at both time points in diabetic *Glut1*^*+/+*^ mice (Fig 3; 2wk a-wave: F(3,210)=20.76, p<0.0001; 4wk a-wave: F(3,195)=40.04, p<0.0001; 2wk b-wave: F(3,420)=59.65, p<0.0001; 4wk b-wave: F(3,390)=112.5, p<0.0001). Representative strobe flash waveforms generated by a 1.4 log cd.s/m^2^ stimulus are shown in Fig 3A and 3D. Luminance response functions of the a- and b-wave are presented in Fig 3B-C and E-F. While the ERGs of *Glut1*^*+/+*^ mice were severely affected by diabetes, no significant defects in the a- or b-wave were found in response to any light stimulus in diabetic *Glut1*^*+/−*^ mice at either time point (see Table 1 for full Two-Way ANOVA and Tukey post-hoc analysis).

**Figure 3:**
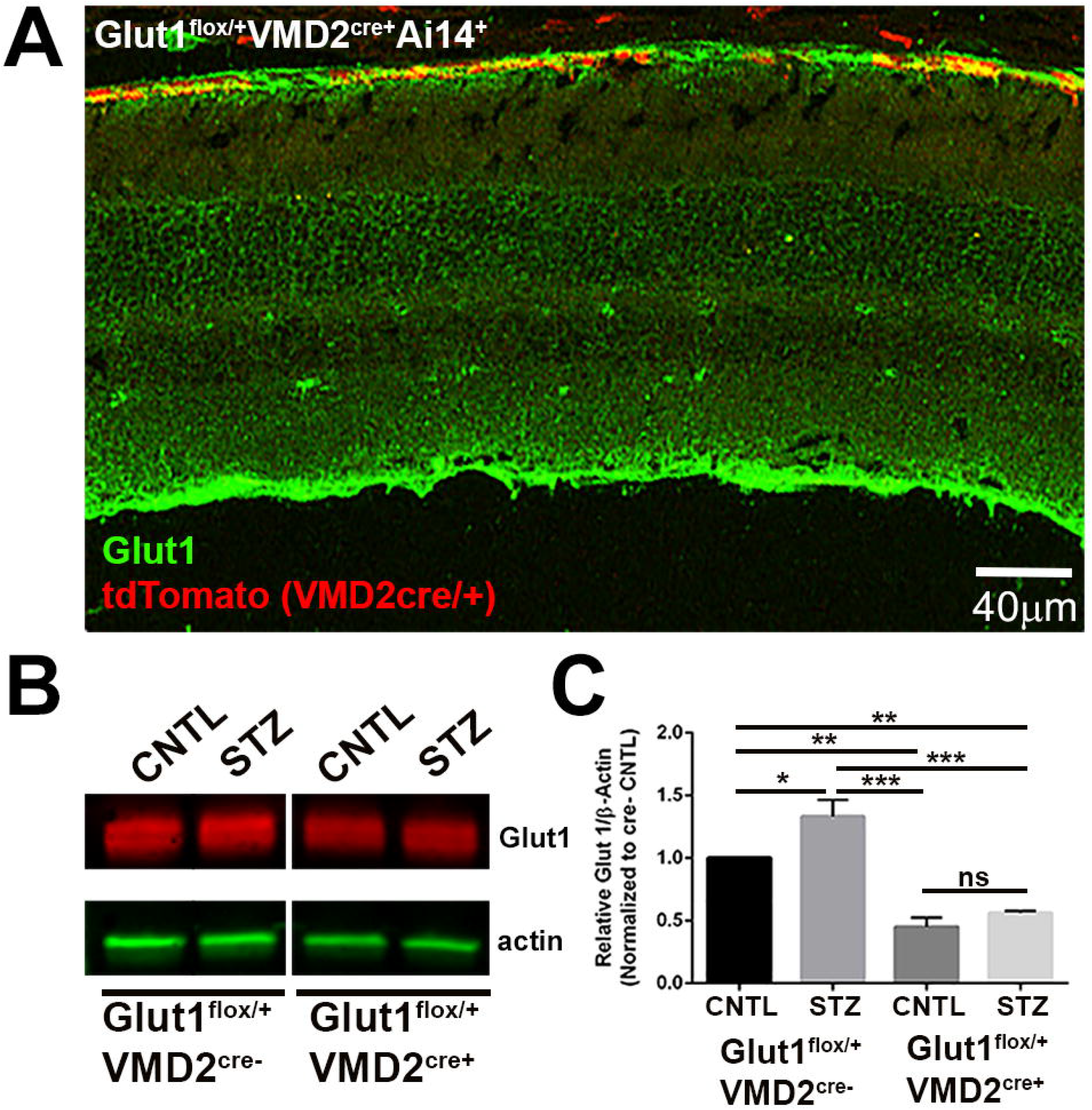
Systemic reduction of Glut1 ameliorates diabetes-induced reductions in ERG component amplitudes. A-C. Representative strobe flash ERG waveform traces evoked in response to a 1.4 log cd.s/m^2^ light stimulus and luminance-response functions for the a-wave and b-wave after 2 weeks of diabetes. D-F. Representative strobe flash ERG waveform traces evoked in response to a 1.4 log cd.s/m^2^ light stimulus and luminance-response functions for the a-wave and b-wave after 4 weeks of diabetes. Amplitude of the a-wave was measured at 8.3msec following the flash stimulus. Amplitude of the b-wave was measured by summing the amplitude of the a-wave with the peak of the response following the oscillatory potentials (≥40 msec). G. Representative traces of filtered oscillatory potentials from strobe flash ERGs evoked by a 1.4 log cd.s/m^2^ flash stimulus at 2- and 4-weeks of diabetes. H-I. Average amplitude of OP1-3 at 2 and 4 weeks of diabetes. Amplitude was measured from the minimum of the preceding trough to the peak of the potential. J. Representative light-adapted waveform traces generated by a 1.4 log cd.s/m^2^ flash stimulus. Light-adapted response amplitudes were calculated by summing the peak of the waveform with the amplitude at 8.3 msec. K-L. Average amplitude of the light-adapted response at 2 and 4 weeks of diabetes. M. Representative waveforms induced by a 5cd/m^2^ white stimulus for 10 seconds. N-O. Average amplitude of the c-wave at 2 and 4 weeks of diabetes. Amplitude of the c-wave was determined by subtracting the average baseline amplitude from the maximal response following the b-wave. All graphs depict mean amplitude ± SEM for each flash stimulus except for the c-wave, which depicts mean ± SD. n≥3 in each group. *p≤0.05; **p≤0.001; ***p≤0.0001. Table 3-1 presents latency times for OP1-3 in response to the 1.4 log cd.s/m^2^ flash stimulus at 2 and 4 weeks of diabetes.

Oscillatory potentials (OPs) were filtered from the 1.4 log cd.s/m^2^ waveform traces at each time point for analysis of this characteristic defect commonly found in diabetic rodents and patients even after only short durations of diabetes (Bresnick et al., 1984; Bresnick and Palta, 1987; Pardue et al., 2014). While no differences in latency of the OPs based on genotype or diabetes status were found (Table 3-1), we identified a significant reduction in OP amplitudes in diabetic *Glut1*^*+/+*^ mice which was not present in diabetic *Glut1*^*+/−*^ mice (Fig 3G-I; 2wk OP1: F(3,28)=7.353, p=0.0009; 2wk OP2: F(3,28)=2.691, p=0.0653; 2wk OP3: F(3,28)=2.990, p=0.0478; 4wk OP1: F(3,31)=6.530, p=0.0015; 4wk OP3: F(3,31)=5.322, p=0.0045). Therefore, in addition to preventing reductions in the a- and b-wave amplitude, systemic lowering of Glut1 also prevented OP amplitude defects. The light-adapted ERG response was also measured at 2- and 4 weeks of diabetes and revealed that diabetic *Glut1*^*+/−*^ mice did not exhibit significant defects in this parameter either (Fig 3J-L; 2wk: F(3, 252)=13.76, p<0.0001; 4wk: F(3, 234)=29.16, p<0.0001). Finally, the RPE-dependent c-wave amplitude was measured in each cohort of mice at both time points. The RPE dependent response in patients is observed with the electro-oculogram. This waveform component is sensitive to glucose and altered in diabetic patients with and without retinopathy (Schneck et al., 2008). Representative c-wave tracings from mice at 2- and 4 weeks of diabetes are shown in Figure 3M. Diabetes significantly reduced the c-wave of *Glut1*^*+/+*^ mice while diabetic *Glut1*^*+/−*^ mice exhibited a less profound defect (Fig 3N-O; 2wk: F(3,60)=15.67, p<0.0001; 4wk: F(3,53)=8.232, p=0.0001).

Retinal inflammation and oxidative stress are characteristic pathological features associated with early states of diabetes, reflecting the overproduction of superoxide and reactive oxygen species (Baynes, 1991; Du et al., 2003; Al-Kharashi, 2018). As such, we investigated if systemic reduction of Glut1 and mitigated retinal polyol accumulation also prevented these markers of DR. Superoxide production was determined by staining fresh frozen retinal sections with dihydroethidium (DHE, Fig 4A). Quantification of total corrected cell fluorescence identified a two-fold increase in retinal superoxide in *Glut1*^*+/+*^ mice while there was no increase in DHE in *Glut1*^*+/−*^ mice (Fig 4B; F(3, 12)=30.73, p<0.0001). Furthermore, quantitative PCR of oxidative stress and inflammation molecules, Nos2 (Fig 4C; F(3,12)=13.95, p=0.0003) and Cox2 (Fig 4D; F(3,8)=5.924, p=0.0198); inflammatory cytokines, IL1-β (Fig4E; F(3,8)=34.22, p<0.0001) and TNF-α (Fig 4F; F(3,8)=52.93, p<0.0001); and angiogenic molecules, VEGF (Fig 4G; F(3,8)=19.00, p=0.0005) and HIF1α (Fig4H; F(3,8)=13.57, p=0.0017) from CNTL and STZ retinas of each cohort of mice revealed that systemic reduction of Glut1 was sufficient to prevent the diabetes-induced increase in each of these molecules.

**Figure 4.**
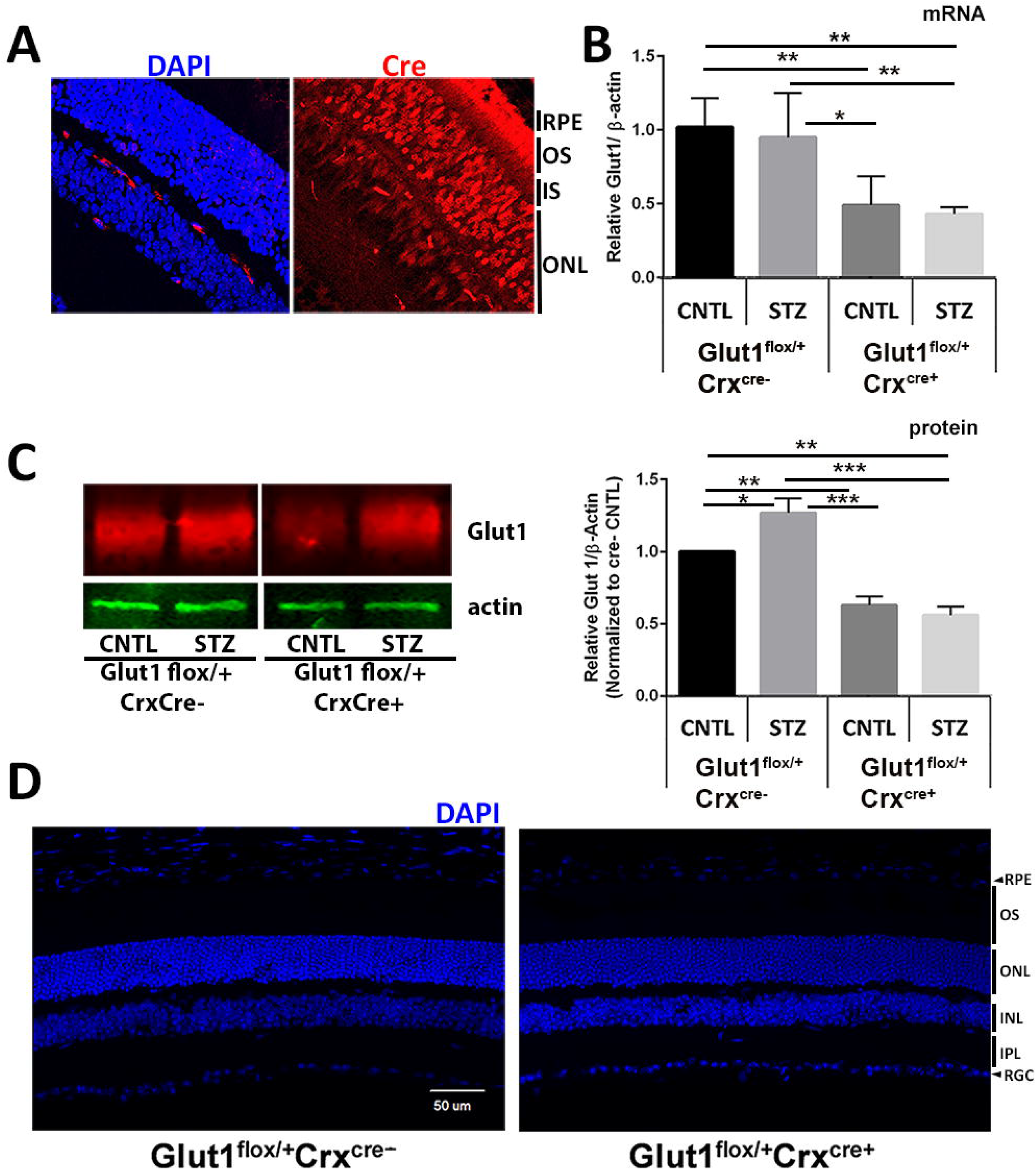
Systemic reduction of Glut1 prevents early elevations in retinal oxidative stress and inflammation. A. Photomicrographs of fresh frozen retinal cryosections from mice at 4 weeks of diabetes probed with dihydroethidium (red). Scale bar = 50 µm. B. Quantification of corrected total cell fluorescence. Three separate images from at least 4 animals of each group were analyzed. Graph represents mean ± SD. C-H. Quantification of oxidative stress molecules and inflammatory cytokines at 4 weeks of diabetes. Graphs represent mean ± SEM. n≥3 for each group. *p≤0.05; **p≤0.001; ***p≤0.0001.

### Reduction of Glut1 in the RPE does not protect against early markers of DR

Our results indicate that systemic reduction of Glut1 protects against multiple early features of DR. To better understand this protection, we next determined if reduction of Glut1 in specific cell types would confer a similar protection. We first examined the RPE, where Glut1 is expressed on both the apical and basal membranes (Kumagai et al., 1994). RPE-specific Glut1 conditional knockdown (CKD) mice were generated by crossing the *VMD2*^*Cre/+*^ strain with the *Glut1*^*flox*^ strain (VMD2 Glut1-CKD). Due to recombination with only one *Glut1*^*flox*^ allele, *VMD2*^*Cre/+*^*Glut1*^*flox/+*^ mice effectively recapitulate the *Glut1*^*+/−*^ phenotype except that Glut1 is reduced by 50% only within the RPE. Figure 5-1A demonstrates *VMD2*^*Cre/+*^-mediated recombination with tdTomato. Nearly all of the RPE cells exhibited tdTomato expression (red) and still retain some Glut1 (green). In comparison to *Glut1*^*flox/flox*^*VMD2*^*Cre/+*^ (Glut1_m_) mice (Swarup et al., 2019), which had moderate levels of patchy Cre expression and induced the complete loss of Glut1 in only about 50% of RPE cells, the VMD2 Glut1-CKD model had higher levels of Cre expression throughout the RPE so that Glut1 was reduced by 50% in most cells due to the deletion of a single *Glut1*^*flox*^ allele. This ensured normal retinal and RPE function (demonstrated in Fig 6). Figure 5-1B-C illustrates the 50% reduction of Glut1 in the RPE by Western blot analysis (F(3,11)=26.33, p<0.0001). Like the diabetic *Glut1*^*+/−*^ retinas, diabetic VMD2 Glut1-CKD mice exhibited overt systemic hyperglycemia (Fig 5A; F(3,12)=24, p<0.0001), and a trend toward higher retinal glucose levels in the diabetic mice, but no significant differences were found between any groups (Fig 5B; F(3,15)=3.5, p=0.0515). However, high levels of retinal sorbitol (Fig 5C; F(3,12)=30, p<0.0001) remained in both diabetic groups. Although multiple comparison differences did not reach significance, elevated levels of fructose were also observed (Fig 5D; F(3,12)=4.2, p=0.0302). Cytosolic NADH/NAD ratio, as identified by lactate:pyruvate ratios, was also unchanged between groups (Fig 5E; F(3,12)=0.83, p=0.5036). To determine whether reduction of Glut1 in the RPE normalized retinal function despite the presence of polyol accumulation, ERGs were performed on diabetic and non-diabetic VMD2 Glut1-CKD and littermate control mice. In line with a role for glucose and glucose metabolites affecting retinal function, both genotypes exhibited reduced ERG waveform components at 4 weeks of diabetes. Figure 6A depicts representative waveform traces evoked by a 1.4 log cd.s/m^2^ stimulus flash from each group of mice. The luminance-response functions for the a- and b-wave are shown in Fig 6B-C. Neither waveform component was rescued by the reduction of Glut1 only in the RPE (a-wave: F(3, 120)=12.75, p<0.0001; b-wave: F(3, 200)=27.25, p<0.0001). Amplitude of filtered OPs (Fig 6D-E) and c-wave amplitudes (Fig 6F-G; F(3,17)=8.042, p=0.0015) were similarly, reduced by equivalent amounts in diabetic wildtype and VMD2 Glut1-CKD mice. These data indicate that reduction of Glut1 in the RPE is not sufficient to prevent retinal polyol accumulation or retinal dysfunction which were both abrogated in diabetic *Glut1*^*+/−*^ mice.

**Figure 5:**
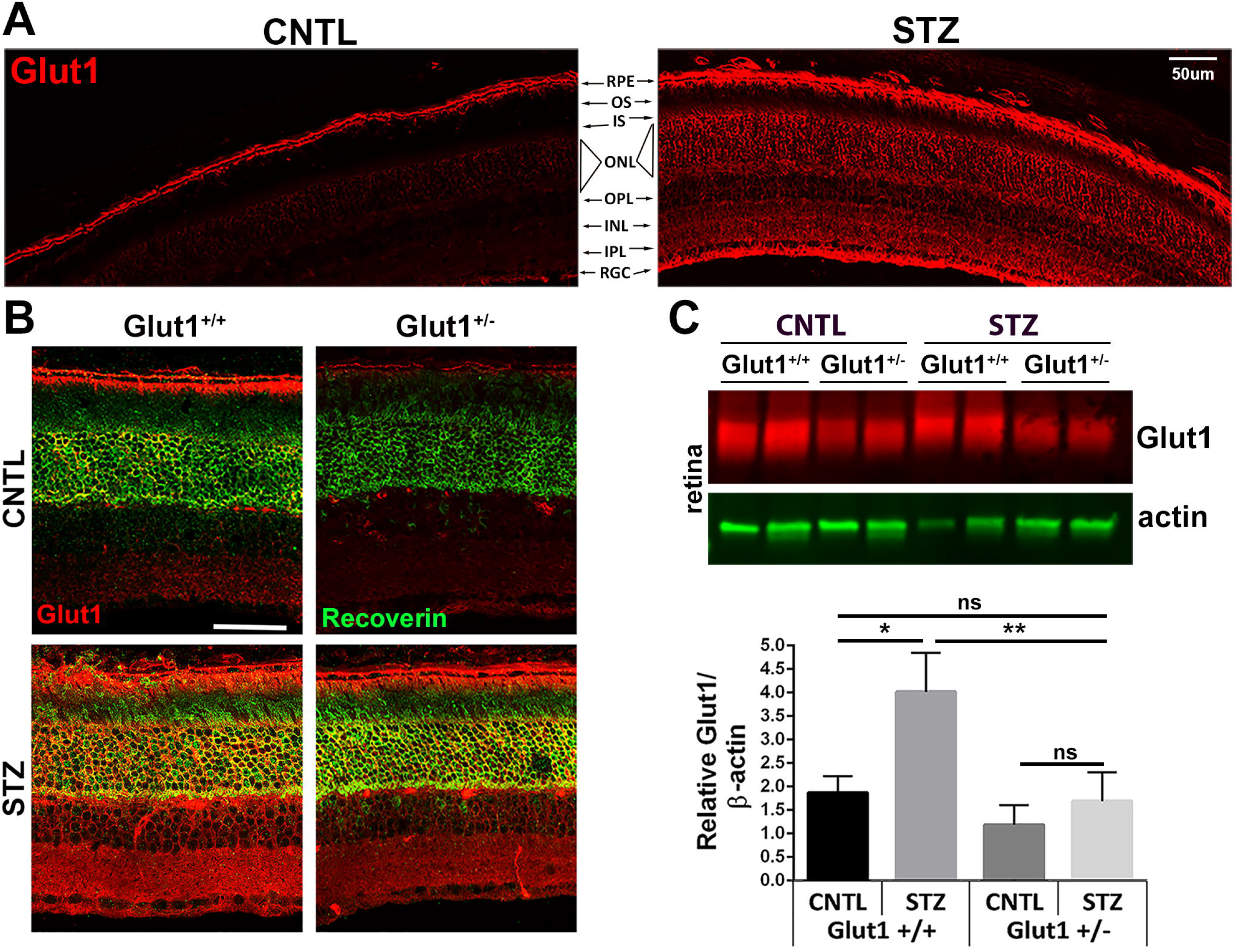
Reduction of Glut1 in the RPE does not mitigate elevations in retinal sorbitol. A. At 4 weeks of diabetes, mice were fasted for ≥7 hours prior to analysis of blood glucose levels with a One-touch Ultra glucometer. B-E. Retinas from fasted VMD2 Glut1-CKD mice were dissected and analyzed by GC/MS after 4 weeks of diabetes. Relative quantities of glucose (B), sorbitol (C) and fructose (D) were normalized to ^13^C_5_-ribitol for comparison between genotypes. E. Comparison of lactate:pyruvate ratios between genotypes. All graphs represent mean ± SD. n=4 for each group. *p≤0.05; **p≤0.001; ***p≤0.0001. Figure 5-1 illustrates VMD2^Cre/+^ expression and 50% reduction of Glut1 in the RPE.

**Figure 6:**
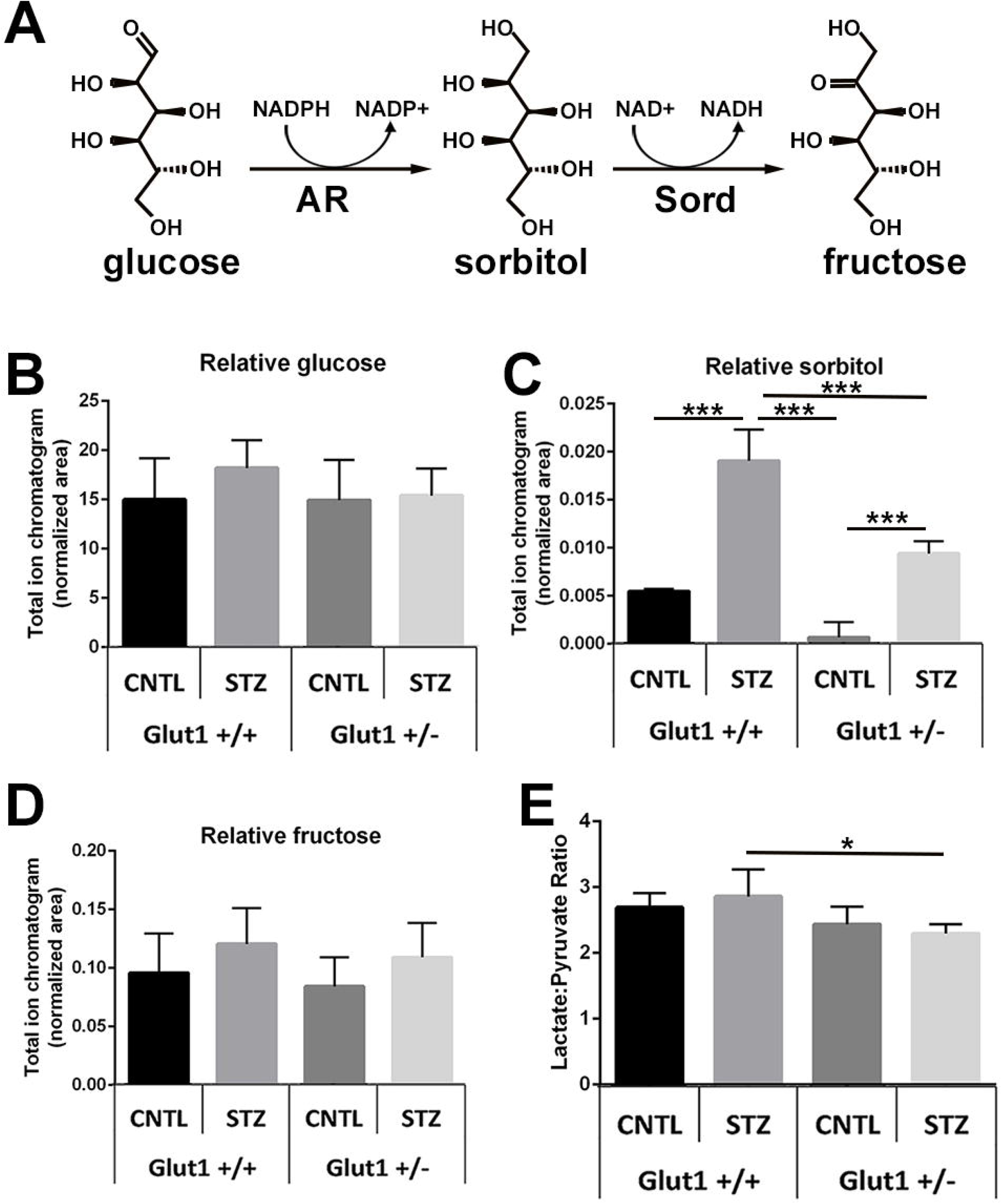
Diabetic mice with reduction of Glut1 in the RPE exhibit similar ERG defects as diabetic controls. A. Representative strobe flash ERG waveform traces from diabetic (STZ) and non-diabetic (CNTL) VMD2 Glut1-CKD mice and littermate controls evoked in response to a 1.4 log cd.s/m^2^ light stimulus. B. Luminance-response function for the a-wave after 4 weeks of diabetes. Amplitude of the a-wave was measured at 8.3msec following the flash stimulus. C. Luminance-response function for the b-wave after 4 weeks of diabetes. Amplitude of the b-wave was measured by summing the amplitude of the a-wave with the peak of the response following the oscillatory potentials (≥40msec). D. Representative traces of filtered oscillatory potentials from strobe flash ERGs evoked by a 1.4 log cd.s/m^2^ flash stimulus at 4-weeks of diabetes. E. Mean amplitude of OP1-3 and the summed OP amplitudes 4 weeks of diabetes. Amplitude of each oscillatory potential was measured from the minimum of the preceding trough to the peak of the potential. F. Representative waveforms induced by a 5cd/m^2^ white stimulus for 10 seconds. Amplitude of the c-wave was determined by subtracting the average baseline amplitude from the maximal response following the b-wave. G. Average amplitude of the c-wave at 4 weeks of diabetes. All graphs depict mean amplitude ± SEM for each flash stimulus. n≥4 in each group. *p≤0.05; **p≤0.001; ***p≤0.0001.

### Reduction of Glut1 in retinal neurons reduces polyol accumulation and prevents retinal dysfunction and markers of inflammation/oxidative stress

We next used the *Crx*^*Cre/+*^ transgenic strain to drive Cre expression in retinal neurons. As with the VMD2 strain, *Glut1*^*flox/+*^ mice were bred with *Crx*^*Cre/+*^ mice to create the retina specific Crx Glut1-CKD mouse that expresses a single *Slc2a1* allele in all the cells of the Crx lineage. Crx is expressed in photoreceptor progenitors beginning at E12.5 and Crx-mediated recombination occurs in rod and cone photoreceptors, bipolar cells and amacrine cells (Furukawa et al., 1997; Furukawa et al., 2002; Hennig et al., 2008). Fig 7-1A illustrates the pattern of *Crx*^*Cre*^ mediated recombination within retinal neurons, but not the RPE. Decreased levels of *Slc2a1* mRNA and Glut1 protein in retinas of Crx Glut1-CKD mice were confirmed by qPCR (Fig 7-1B; F(3,18)=9.573, p=0.0005) and western blotting (Fig 7-1C; F(3,15)=25.29, p<0.0001), respectively. Like *Glut1*^*+/−*^ mice, non-diabetic Crx Glut1-CKD mice also exhibited normal retinal morphology (Fig 7-1D) and function (Fig 8).

**Figure 7:**
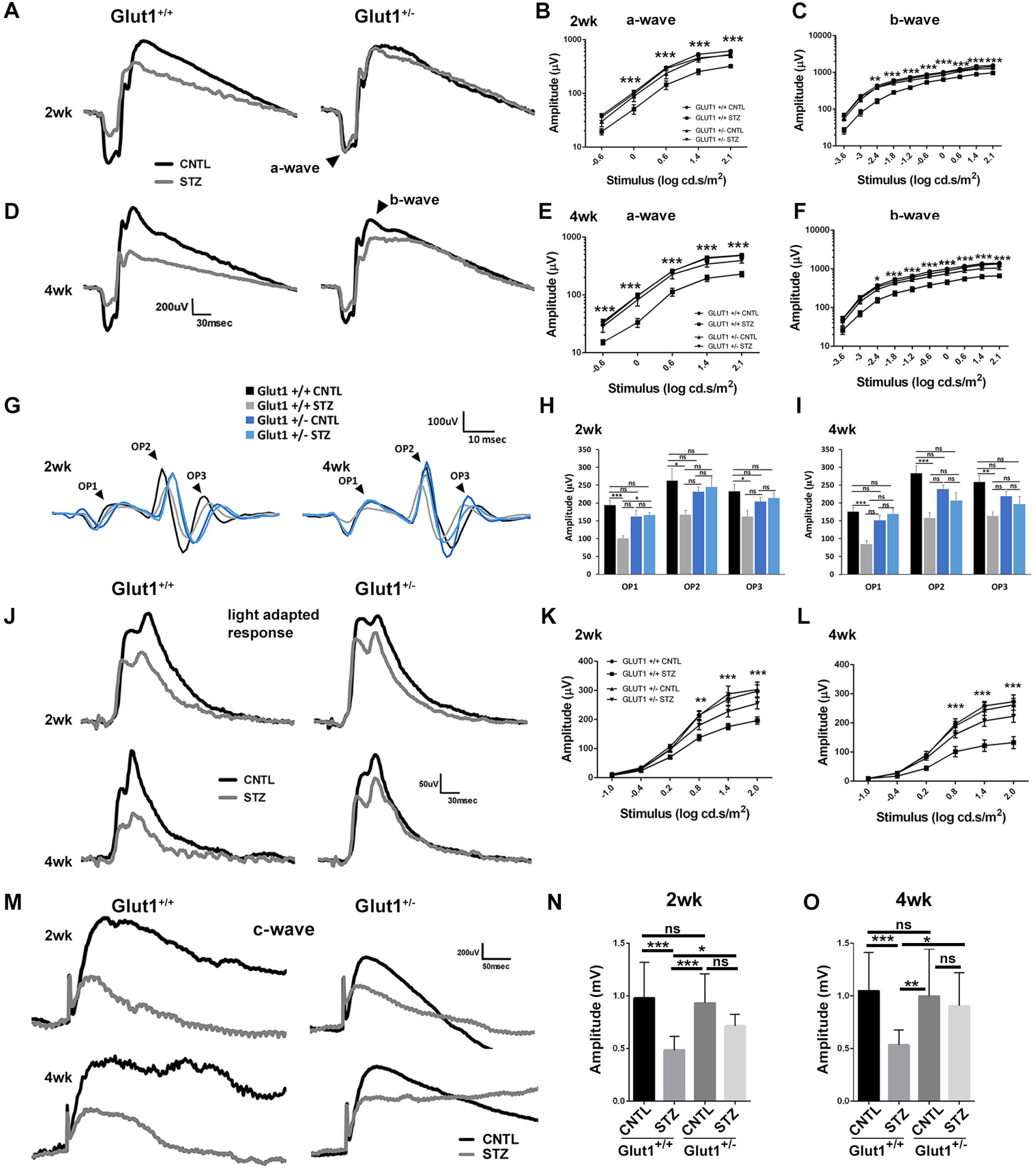
Reduction of Glut1 in the retina ameliorates retinal polyol accumulation associated with diabetes. A. At 4 weeks of diabetes, mice were fasted for ≥7 hours prior to analysis of blood glucose levels with a One-touch Ultra glucometer. B-E. Retinas from fasted Crx Glut1-CKD mice were dissected and analyzed by GC/MS. Relative quantities of glucose (B), sorbitol (C) and fructose (D) were normalized to ^13^C_5_-ribitol for comparison between genotypes. E. Comparison of lactate:pyruvate ratios between genotypes. All graphs represent mean ± SD. n=5 for each group. *p≤0.05; **p≤0.001; ***p≤0.0001. Figure 7-1 illustrates Crx^Cre/+^ expression and 50% reduction of Glut1 in retinal neurons.

**Figure 8.**
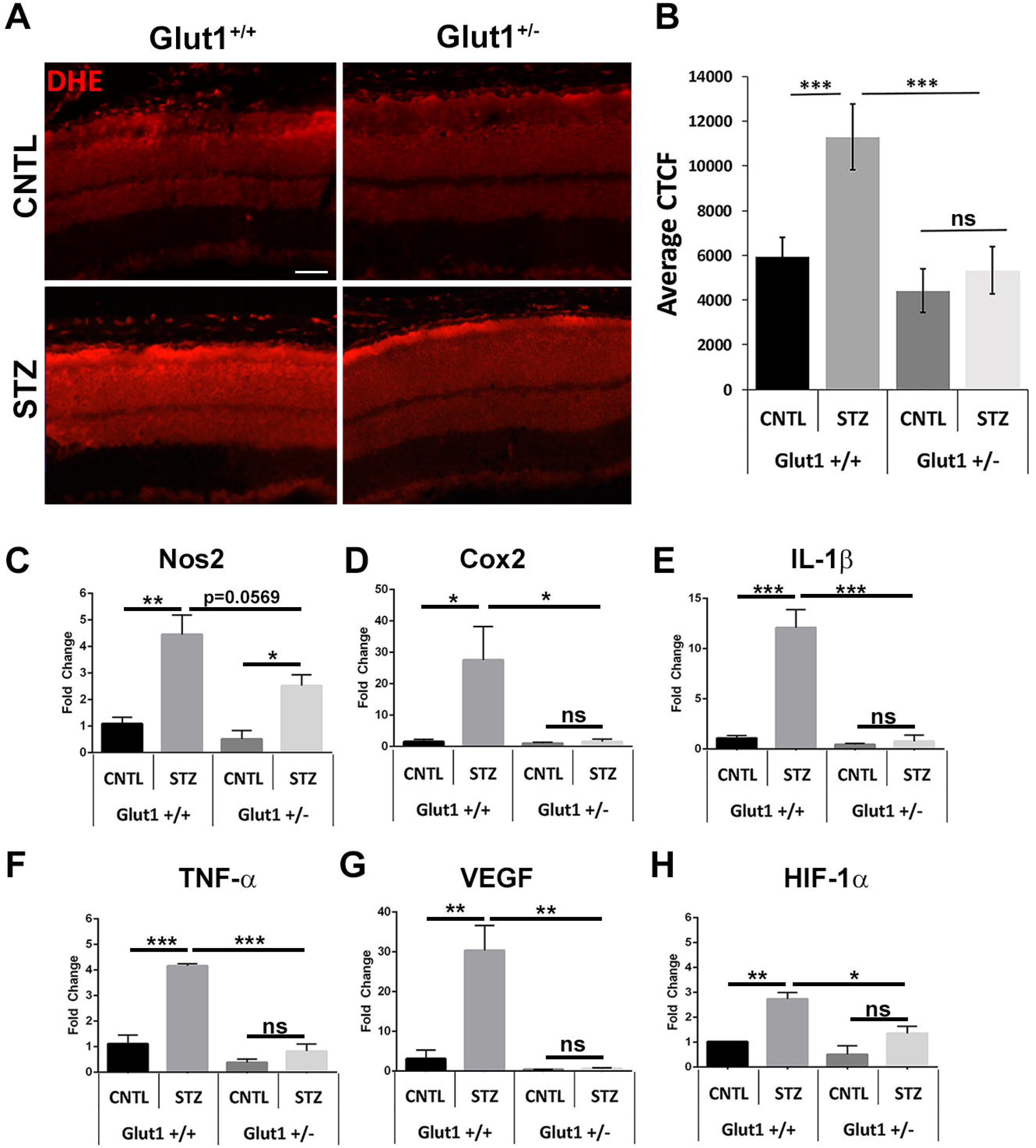
Diabetic mice with reduction of Glut1 in the retina exhibit no ERG defects. A. Representative strobe flash ERG waveform traces from diabetic and non-diabetic Crx Glut1-CKD mice and littermate controls evoked in response to a 1.4 log cd.s/m^2^ light stimulus. B. Luminance-response function for the a-wave after 4 weeks of diabetes. Amplitude of the a-wave was measured at 8.3msec following the flash stimulus. C. Luminance-response function for the b-wave after 4 weeks of diabetes. Amplitude of the b-wave was measured by summing the amplitude of the a-wave with the peak of the response following the oscillatory potentials (≥40 msec). D. Representative traces of filtered oscillatory potentials from strobe flash ERGs evoked by a 1.4 log cd.s/m^2^ flash stimulus at 4-weeks of diabetes. E. Average amplitude of OP1-3 and the summed OP amplitudes 4 weeks of diabetes. Amplitude of each oscillatory potential was measured from the minimum of the preceding trough to the peak of the potential. F. Representative waveforms induced by a 5 cd/m^2^ white stimulus for 10 seconds. Amplitude of the c-wave was determined by subtracting the average baseline amplitude from the maximal response following the b-wave. G. Average amplitude of the c-wave at 4 weeks of diabetes. All graphs depict mean amplitude ± SEM for each flash stimulus. n≥3 in each group. *p≤0.05; **p≤0.001; ***p≤0.0001.

Analysis of blood and retinal glucose levels revealed that reduction of Glut1 only in the retina did not affect systemic glucose levels (Fig 7A; F(3,16)=31, p<0.0001), but slightly reduced retinal glucose content, which was only significant in comparison with the wildtype diabetic retina (Fig 7B; F(3,16)=4.4, p=0.0196). There were no differences in systemic or retinal glucose levels between diabetic mice based on genotype, however. Importantly, diabetic Crx Glut1-CKD mice displayed a significant reduction in retinal sorbitol levels in comparison to wildtype diabetics (Fig 7C; F(3,15)=34, p<0.0001). Similar to the glucose content, fructose levels in the non-diabetic Crx Glut1-CKD mouse were slightly lower, but only in comparison to the wildtype diabetic (Fig 7D; F(3,16)=4.6, p=0.0160). No differences were found in lactate:pyruvate ratios. These findings reveal that reduction of Glut1 only within retinal neurons can recapitulate the mitigation of retinal polyol accumulation found in diabetic *Glut1*^+/−^ mice. Importantly, although reduction of Glut1 only in retinal neurons did not lead to a complete normalization of retinal sorbitol levels, the change was correlated with a full normalization of retinal function (Fig 8). ERGs were recorded following 4 weeks of diabetes in Crx Glut1-CKD and littermate controls. Notably, diabetic Crx Glut1-CKD mice displayed normal a- and b-wave amplitudes at all light intensities (Fig 8A-C; a-wave: F(3,142)=24.74, p<0.0001; b-wave: F(3,233)=42.71, p<0.0001). Figure 8A illustrates waveform traces from a 1.9 log cd.s/m^2^ flash stimulus and in Fig 8B-C, luminance-response functions for these mice clearly depict the significant differences between diabetic wildtype and Crx Glut1-CKD mice. The amplitude of the oscillatory potentials (Fig 8D-E) and the c-wave (Fig 8F-G; F(3,25)=23.38, p<0.0001) was also normalized in the diabetic Crx Glut1-CKD mouse. To determine if the rescue of the diabetic phenotype extended beyond retinal function, we analyzed expression of oxidative stress molecules and inflammatory cytokines in retinas from each cohort of animals (Fig 9). While diabetes induced elevations in each molecule/cytokine in wildtype mice, no differences in expression of Nos2 (Fig 9A; F(3,13)=6.710, p=0.0056), TNF-α (Fig 9B; F(3,16)=23.34, p<0.0001), Cox2 (Fig 9C; F(3, 13)=9.133, p=0.0016), or IL-1β (Fig 9D; F(3, 14)=10.23, p=0.0008) were observed between non-diabetic wildtype and diabetic Crx Glut1-CKD cohorts. These findings demonstrate that a small modulation of Glut1 levels only within retinal neurons has significant effects on the development of early characteristic hallmarks of DR.

**Figure 9:**
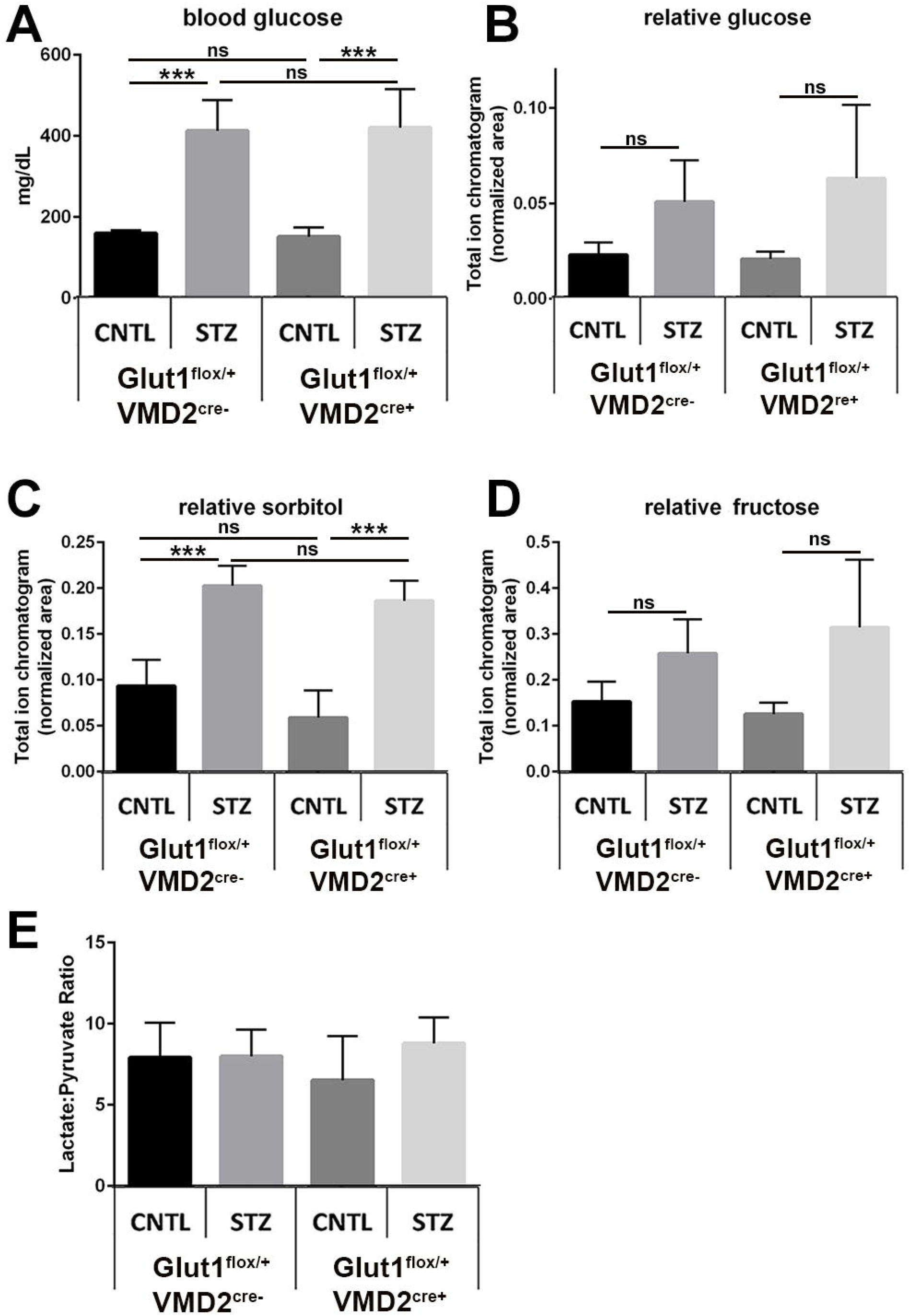
Diabetic mice with reduction of Glut1 in retinal neurons exhibit normalized levels of oxidative stress and inflammatory markers. Expression of Nos2 (A), TNF-a (B), Cox2 (C) and IL-1b (D) in retinas from 4-week diabetic mice were measured by quantitative PCR. Graphs present mean ± SD. n≥3 for each group. *p≤0.05; **p≤0.001; ***p≤0.0001.

## Discussion

Almost 20 years ago a unifying theory for the etiology of DR was proposed. It postulated that hyperglycemia-induced production of free oxygen radicals was the basis for glucose damage in the retina (Nishikawa et al., 2000; Brownlee, 2001, 2005). Glucose toxicity and oxidative stress lead to retinal vascular damage through multiple downstream mechanisms including activation of protein kinase C, aldose reductase activation, and advanced glycation end product formation (Du et al., 2000). Despite this unifying theory, and a multitude of attempts to intervene in the molecular and biochemical pathways stemming from oxidative stress and hyperglycemia, the goal of preventing DR has not yet been met. Therefore, investigation into the mechanisms of glucose entry into the retina, glucose metabolism, and the production of oxidative damage has continued. We found here that (1) Glut1 is elevated at early stages of DR, (2) systemic reduction of Glut1 is a successful mechanism for the prevention of polyol accumulation, functional defects, and increased markers of inflammation and oxidative stress within the diabetic retina, and (3) reduction of Glut1 suppressed these early hallmarks of DR when it was targeted in retinal neurons, but not in the RPE.

Glut1 is the primary facilitative transporter for the retina and is expressed almost ubiquitously in ocular tissues. It is located on the apical and basal RPE membranes and on the luminal and abluminal membranes of retinal endothelial cells to facilitate glucose flux in the retina. It is also expressed on retinal ganglion cells and is thought to be the only known glucose transporter expressed by photoreceptors (Mantych et al., 1993; Gospe et al., 2010). We found that retinal Glut1 is elevated in early DR, and document this by immunohistochemistry and western blotting. Furthermore, quantitative proteomics revealed a significant increase in retinal Glut1 levels in both male and female diabetic C57Bl/6J mice at 3 weeks of diabetes [Females: average linear ratio = 2.519; average Ln Ratio = 0.924; moderated p-value=6.1×10^−3^**; moderated adjusted p-value=6.1×10^−3^**; 4/4 mice. Males: average linear ratio = 2.020; average Ln Ratio = 0.703; moderated p-value=1.3×10^−2^**; moderated adjusted p-value=3.3×10^−2^**; 4/4 mice (**elevations≥ 2SD, personal communication from Dr. Bela Anand-Apte, Cole Eye Institute, Cleveland Clinic)]. However, the literature is inconsistent in this respect, with conflicting reports demonstrating no change (Kumagai et al., 1994; Antonetti et al., 1998), reduced (Badr et al., 2000; Fernandes et al., 2004) and increased (Kumagai et al., 1996; Lu et al., 2013) Glut1 levels in the diabetic retina. Resolution of this divergence is complicated by variability in methodology of animal maintenance (insulin treatment/frequency/concentration), timing of analysis, and the tissue target for analysis. While the basis for the differences is not entirely clear, and unlikely to be resolved, our findings unequivocally show that Glut1 was elevated in the diabetic retina at early time points, which correlated with initial indices of DR.

The profound protection against ERG defects and early markers of oxidative stress/inflammation that we found in *Glut1*^*+/−*^ mice agrees with previous studies demonstrating a role for Glut1 in development of DR (Lu et al., 2013; You et al., 2017; You et al., 2018). Our work is novel in that we systemically reduced Glut1 by a genetic approach, correlated it with normalization of polyol accumulation and determined that reduction of Glut1 in the RPE is not protective. It is important to note that although glucose is required for function and survival of photoreceptors, and these cells undergo degeneration in mice with ≥50% reduction of Glut1 in the RPE (Swarup et al., 2019), the retina is resilient to ≤50% reduction of Glut1 in the RPE. We demonstrated here by ERG and histological analysis that no changes in retinal function or morphology of the retina are found in *Glut1*^*+/−*^ mice, a model of *Slc2a1* haploinsufficiency and Glut1 deficiency syndrome (Wang et al., 2006). Although neuroinflammation and microvascular changes occur in the brain of *Glut1*^*+/−*^ mice (Tang et al., 2017; Tang et al., 2019)), *Glut1*^*+/−*^retinas are normal. Likewise, inactivation of one *Slc2a1* allele in the retina (Crx Glut1-CKD) or the RPE (VMD2 Glut1-CKD) also maintained normal electroretinography and histology, with lower levels of Glut1 in these cell types but no compensation by other glucose transporters (RT-qPCR; other glucose transporters are undetected in the retina by western blot or immunohistochemistry). Therefore, modulation of Glut1 by small amounts in the retina is a feasible strategy for protection against early hallmarks of DR.

STZ induced blood glucose levels to 2-3x of that found in non-diabetic mice (3.1x for *Glut1*^*+/+*^, 3.2x for *Glut1*^*+/−*^; 2.6 for VMD2 controls, 2.8 for VMD2 Glut1-CKDs; 2.3 for Crx controls, 2.4 for Crx Glut1-CKDs). However, *retinal* glucose levels were not significantly different between diabetic and non-diabetic mice of any genotype. Instead, significant differences were found in levels of glucose metabolites of the polyol pathway. Importantly, mice were fasted for ≥7 hours prior to retinal dissections. This enabled a clear evaluation of retinal glucose levels and metabolite accumulation. While glucose can readily be transported out of the retina via Glut1 localized on retinal endothelial cells and the RPE, sorbitol cannot be exported from the retina (Jedziniak et al., 1981). Therefore, we purport that the primary effector in DR is not glucose itself, but sorbitol, which accumulates to lead to increased osmolarity as well as oxidative stress.

Systemic or retina-specific reduction of Glut1 was correlated with lower sorbitol accumulation, and more importantly, with complete normalization of ERG defects and oxidative stress in diabetic mice. Sorbitol induces hyperosmolarity and oxidative stress due to the biochemical processes underlying its production and breakdown. When aldose reductase turns glucose into sorbitol, NADPH is converted to NADP^+^. In hyperglycemia, the requirement for breakdown of excess glucose via the polyol pathway depletes NADPH, which is critical for glutathione to scavenge free radicals and the *de novo* synthesis of fatty acids, nucleotides, steroids and cholesterol. Net formation of fructose from glucose via sorbitol dehydrogenase also results in the breakdown of NADPH and formation of NADH. Even the smallest change in NADH levels is deleterious to the cells. NAD+/NADH ratio in the cell is around 600:1, and a very small change in NADH can decrease this ratio drastically (Ido, 2007), inducing pseudohypoxia and contributing to DR (Williamson et al., 1993). Increased NADPH/NADP^+^ (Varma, 1974) and decreased NAD^+^/NADH ratios (Obrosova et al., 2001) have been reported in diabetic rat lens. Although lactate:pyruvate ratios were largely unchanged in our mice, the prevention of polyol accumulation may have directly led to the prevention of oxidative stress.

Perhaps the most important finding from our work was that VMD2 Glut1-CKD mice did not exhibit altered sorbitol levels. Indeed, it was surprising that reduction of Glut1 in the RPE was not associated with lower retinal polyol accumulation or normalized ERGs. However, because glucose is rapidly converted to sorbitol, it was likely that the level of Glut1 remaining on the RPE was too high to sufficiently lower glucose flux into the retina and affect metabolism. Instead, the reduction in glucose entry into retinal neurons was required for the successful normalization of characteristic DR pathologies. Thus, targeting sorbitol or identifying mechanisms to reduce Glut1 in neurons is likely to be key to preventing DR. Crx-mediated recombination occurs in retinal progenitors at E12.5 and results in recombination in photoreceptors, but also bipolar cells and amacrine cells (Hennig et al., 2008). Photoreceptors are the most highly metabolic cells in the body, and maintenance of the dark current is a considerable energy sink for the retina (Okawa et al., 2008). Because of this, we propose that reduced photoreceptor-mediated glucose metabolism accounts for the reduction in sorbitol accumulation and the oxidative stress. Interestingly, diabetic *Gnat1*^*−/−*^ mice exhibit significantly reduced leukostasis and cytokine production (Liu et al., 2019). Hurley and colleagues (Du et al., 2016) demonstrated that phototransduction influences metabolic flux and *Gnat1*^*−/−*^ mice no longer display light-evoked metabolic flux. Thus, the protective effects seen in the diabetic *Gnat1*^*−/−*^ mouse could also be due to reduced sorbitol accumulation.

Beyond sorbitol, retinal endothelial cells significantly contribute to DR pathology downstream of inflammation (Fu et al., 2016; Sorrentino et al., 2018). Because systemic or neuron-specific reduction of Glut1 abrogated cytokine expression, we postulate that leukocytes should not be activated to mediate the inflammatory processes leading to retinal endothelial cell death and angiogenesis. Kern and colleagues suggested that photoreceptors communicate with leukocytes via cytokines to propagate inflammation in the retina and kill retinal endothelial cells (Liu et al., 2016; Tonade et al., 2016; Tonade et al., 2017; Liu et al., 2019). Although we focused here on the early stages of retinal dysfunction associated with DR, we propose that reduction of Glut1 in photoreceptors will also prevent REC loss and later features of DR. Determining whether reduction of Glut1 in RECs prevents DR pathology will be informative in further discerning if limiting glucose entry to the retina, or reducing the rate of glucose metabolism by photoreceptors is the source of pathogenicity. Long–term reduction of polyol accumulation and production of free oxygen radicals will still be the key to prevention and treatment of DR.

## Acknowledgements

This work was supported by VA Merit Award I01 BX002754-01A2 (ISS), NIH R01 EY024972 (JES) NIH P30 EY025585, Research to Prevent Blindness Challenge Grant, Cleveland Eye Bank Foundation Grant and funds from Cleveland Clinic Foundation. We extend our gratitude to Drs. E. Dale Abel, Joshua Dunaief and Sujata Rao for providing mice and to Dr. Neal Peachey for critical reading of the manuscript and support of these experiments. Matthew Tarchick’s current affiliation is with the Department of Biology, University of Akron, Akron, OH 44325.

---

**Figure 1-1:**
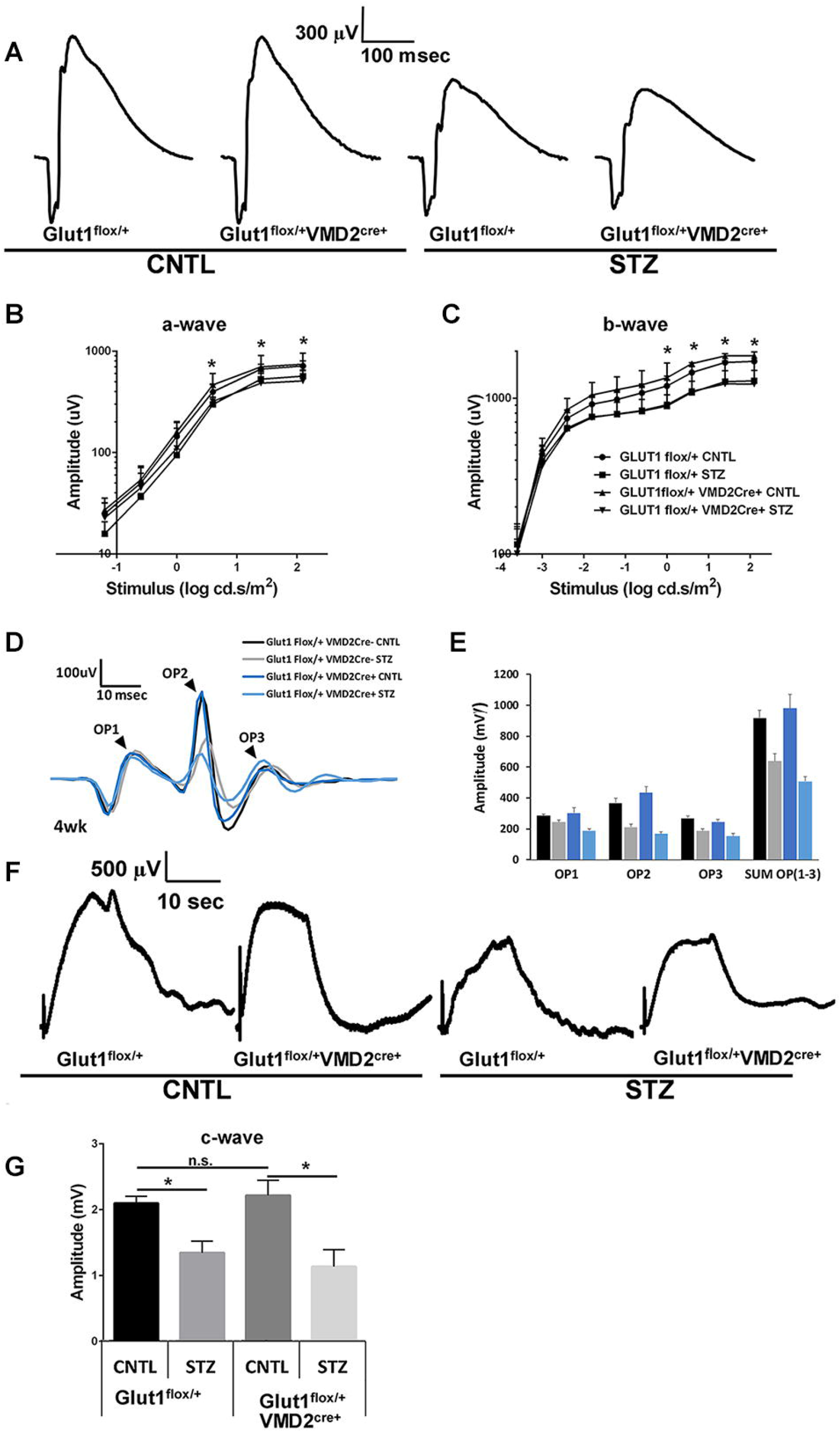
*Glut1*^*+/−*^ mice exhibit normal retinal morphology. A. Representative light photomicrographs of semi-thin plastic sections stained with Toluidine blue O from nondiabetic (CNTL) and diabetic (STZ) mice after 4 weeks of diabetes. Scale bar = 50µm. RPE, retinal pigmented epithelium; OS, outer segments; IS, inner segments; ONL, outer nuclear layer; INL, inner nuclear layer; RGC, retinal ganglion cell layer. B-F. Cell layers were measured from three locations in each image. At least three images per mouse were analyzed.

**Figure 1-2:**
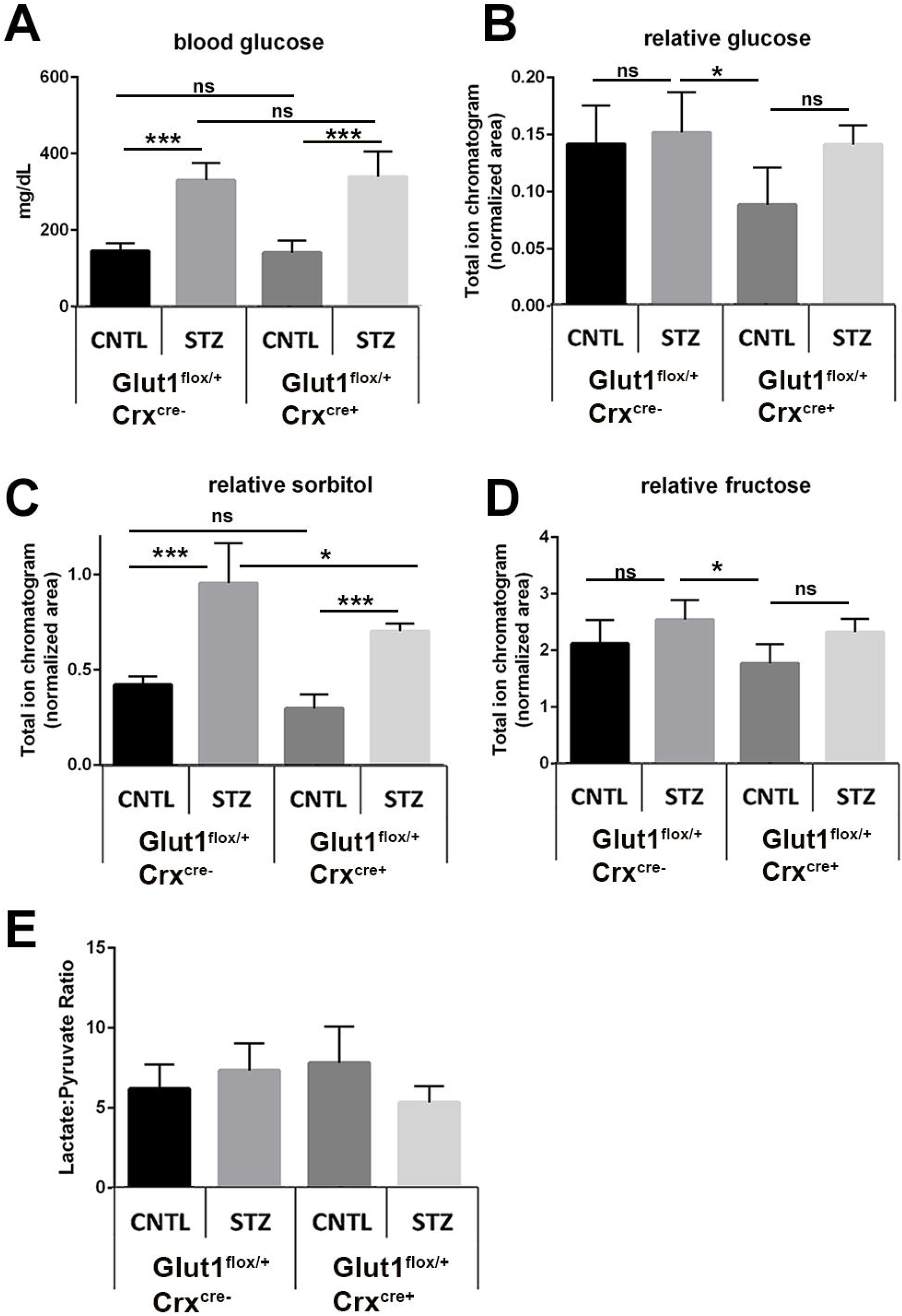
*Glut1*^*+/−*^ mice display normal electroretinography and responses to diabetes. A. Body weight of non-diabetic and diabetic *Glut1*^*+/−*^ and littermate control mice was measured after 4 weeks of diabetes. No differences in weight were identified. B. Mice were fasted for ≥7 hours and blood glucose levels were measured with a One-touch Ultra glucometer. No difference in magnitude of hyperglycemia was observed. C-F. Strobe flash electroretinography was performed on non-diabetic *Glut1*^*+/+*^ and *Glut1*^*+/−*^ mice at 8 weeks of age, prior to induction of diabetes. C. Representative strobe flash ERG waveform traces evoked in response to a 1.4 log cd.s/m^2^ light stimulus. D. Luminance-response functions for the a-wave and b-wave. E. Representative light-adapted ERG waveform traces evoked by a 1.4 log cd.s/m^2^ light stimulus superimposed over the adapting field. F. Luminance-response function for the light-adapted response. No differences in retinal function were found between genotypes.

**Table 1-1:** Diabetes does not alter expression of glucose transporters in the retina. At 4 weeks of diabetes, RNA was extracted from dissected retinas and real-time quantitative PCR was used to analyze expression of glucose transporters in the retina. Relative fold changes in gene expression were determined using the comparative Ct method (2ΔΔCt method).

| Table 1-1: Diabetes does not alter expression of glucose transporters in the retina |  |  |  |  |  |
| --- | --- | --- | --- | --- | --- |
|  | Glut1/*Slc2a1* | Glut3/*Slc2a3* | Glut4/*Slc2a4* | Glut8/*Slc2a8* | Glut12/*Slc2a12* |
| Glut1+/+ CNTL | 1.01 ± 0.08 (4) | 1.01 ± 0.09 (4) | 1.03 ± 0.14 (4) | 1.05 ± 0.22 (3) | 1.06 ± 0.21 (4) |
| Glut1 +/+ STZ | 0.80 ± 0.11 (3) | 0.73 ± 0.21 (3) | 0.97 ± 0.15 (5) | 1.19 ± 0.34 (4) | 0.84 ± 0.21 (3) |
| Glut1+/- CNTL | 0.45 ± 0.07 (4)** | 1.13 ± 0.26 (6) | 1.12 ± 0.13 (8) | 1.25 ± 0.24 (4) | 1.04 ± 0.11 (4) |
| Glut1 +/- STZ | 0.59 ± 0.08 (3)* | 0.81 ± 0.16 (3) | 1.25 ± 0.25 (6) | 0.96 ± 0.21 (3) | 0.86 ± 0.16 (3) |
| Mean fold change ± SEM (n) | | | | | |
| **$p \leq 0.005$, * $p \leq 0.05$ relative to Glut1<sup>+/+</sup> CNTL | | | | | |

**Figure 2-1:**
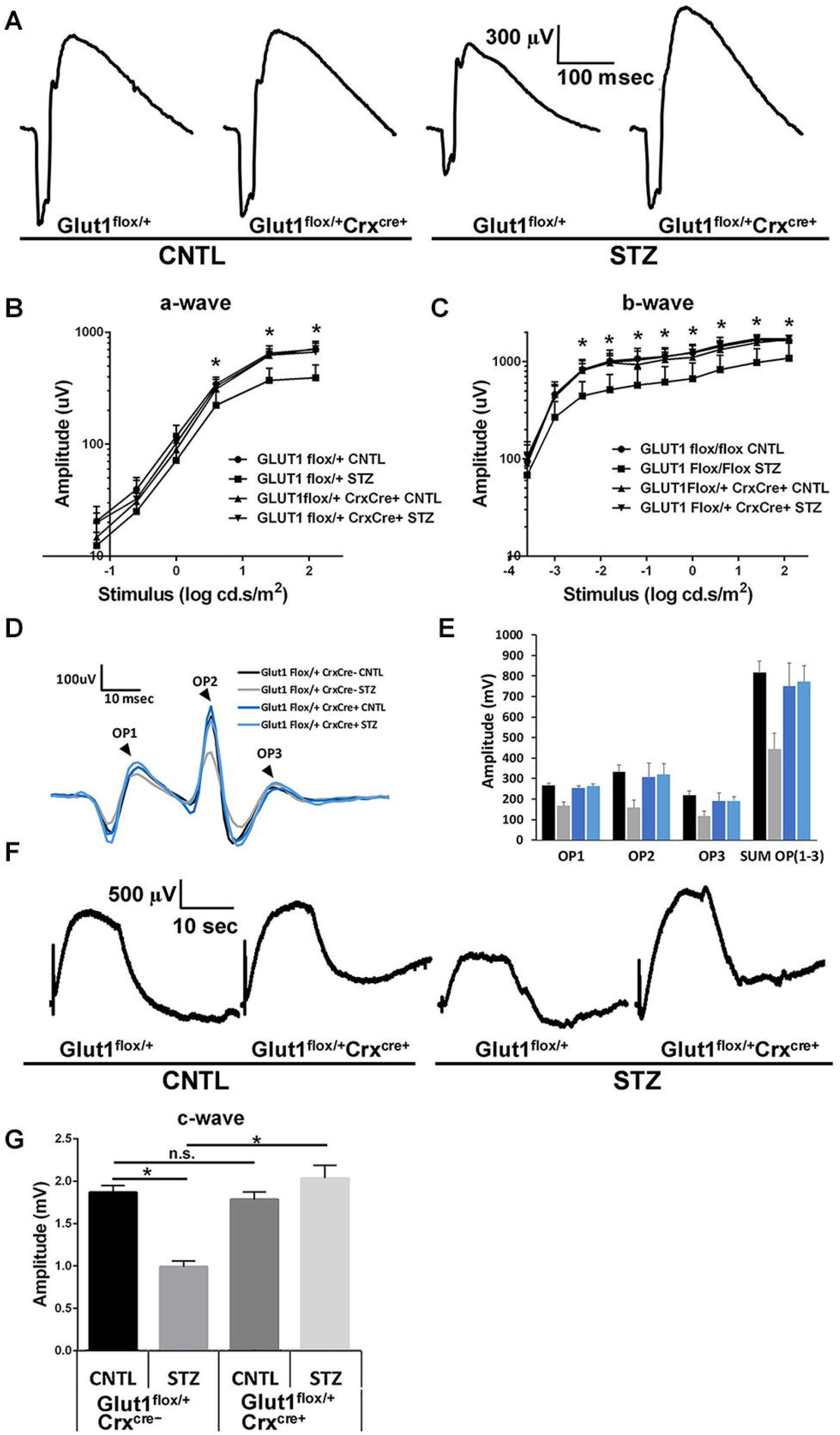
Sorbitol was distinguished from mannitol using retention time. Extracted ion chromatogram m/z 319 of mannitol and sorbitol authentic standards demonstrating baseline separation of these compounds on the GC column.

**Table 3-1:** Diabetes does not alter oscillatory potential latency regardless of Glut1 genotype at early time points. Oscillatory potentials were filtered from strobe flash ERGs evoked by a 1.4 log cd.s/m^2^ flash at 2 and 4 weeks of diabetes. Latency was determined by identifying the time of the peak of each OP wavelet.

| Table 3-1: Oscillatory Potential Latency [Mean msec ± SEM (n)] | | | |
| --- | --- | --- | --- |
| 2wk diabetes | OP1 | OP2 | OP3 |
| Glut1+/+ CNTL | 12.06 ± 0.186 (10) | 27.71 ± 1.081 (10) | 40.19 ± 1.398 (10) |
| Glut1 +/+ STZ | 11.88 ± 0.434 (7) | 24.72 ± 0.566 (7) | 36.73 ± 1.018 (7) |
| Glut1+/- CNTL | 12.65 ± 0.299 (10) | 26.37 ± 0.448 (10) | 38.27 ± 0.644 (10) |
| Glut1 +/- STZ | 12.15 ± 0.564 (5) | 24.46 ± 0.424 (5) | 35.78 ± 0.526 (5) |
| 4wk diabetes |  |  |  |
| Glut1+/+ CNTL | 11.84 ± 0.196 (17) | 24.80 ± 0.353 (17) | 35.23 ± 0.360 (17) |
| Glut1 +/+ STZ | 11.65 ± 0.256 (7) | 23.77 ± 0.509 (7) | 34.71 ± 0.594 (7) |
| Glut1+/- CNTL | 12.00 ± 0.168 (7) | 24.48 ± 0.745 (7) | 36.13 ± 0.809 (7) |
| Glut1 +/- STZ | 11.65 ± 0.340 (4) | 23.71 ± 0.990 (4) | 35.36 ± 1.101 (4) |

**Figure 5-1:VMD2 CKD mice exhibit 50% reduction of Glut1 specifically in the RPE.** A. Representative confocal image demonstrating the *VMD2*^Cre/+^ recombinase-mediated activity in the RPE. Mice with tdTomato (Ai14) expression also have less Glut1. Cre expression is found throughout the RPE but not in the retina. B. Protein levels of Glut1 from RPE tissue isolated from retinas following 4 weeks of diabetes. Retinas were removed from the back of the eye and RPE was isolated in lysis buffer. Glut1 levels were normalized to β-actin. D. Quantitative analysis of Glut1 levels in the RPE. All graphs depict mean ± SEM. n≥4 in each group. *p≤0.05; **p≤0.001; ***p≤0.0001.

**Figure 7-1:**
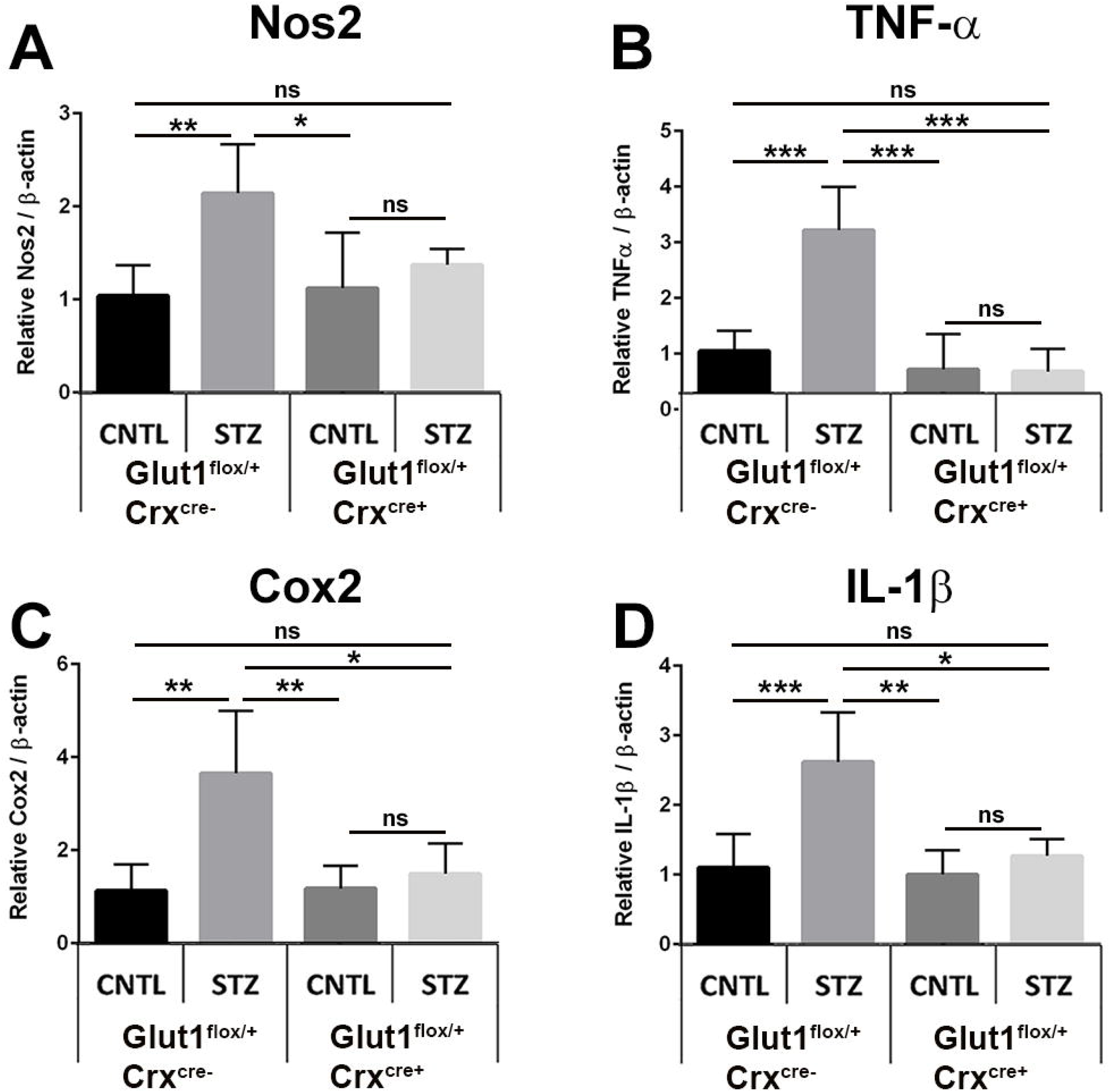
Crx CKD mice exhibit 50% reduction of Glut1 specifically in the retina and normal retinal morphology. A. Representative confocal images depicting *Crx*^Cre/+^ recombinase activity in the retina of adult mice by expression of tdTomato. B. Quantitative PCR demonstrating 50% reduction in Glut1 expression in the retina of *Glut1*^*flox/+*^*Crx*^*cre/+*^mice. C. Protein levels of Glut1 from retinas dissected from *Glut1*^*flox/+*^*Cre*^*Cre/+*^ mice following 4 weeks of diabetes. Left panels depict representative western blots imaged; right graph presents relative Glut1:β-actin levels normalized to the *Glut1*^*flox/+*^*Crx*^*Cre/-*^ control. D. Representative images of retinal cryosections stained with DAPI demonstrating normal retinal morphology. Scale bar = 50 µm. RPE, retinal pigmented epithelium; OS, outer segments; IS, inner segments; ONL, outer nuclear layer; OPL, outer plexiform layer; INL, inner nuclear layer; IPL, inner plexiform layer; RGC, retinal ganglion cell layer. Graphs depict mean ± SEM. n≥3 in each group. *p≤0.05; **p≤0.001; ***p≤0.0001

## Notes

**Conflict of Interest:** The authors declare no competing financial interests.

### Competing Interest Statement

The authors have declared no competing interest.

